# Efficient detection of cryptic pockets on challenging drug targets using MD-free co-vibrational modes

**DOI:** 10.64898/2026.10.07.757394

**Authors:** Tibor Viktor Szalai, Dávid Bajusz, Zoltán Orgován, Péter Ábrányi-Balogh, Márton Gadanecz, Tímea Imre, Emma K. Grant, Jacob T. Bush, István Kolossváry, György M. Keserű

## Abstract

Allosteric and cryptic binding sites provide unique opportunities in drug discovery for achieving selectivity, reducing toxicity and unlocking challenging therapeutical targets. Yet, cryptic pockets are particularly elusive in that they are difficult to capture with experimental methodologies, and therefore often remain undetected. Computational approaches for flexible cryptic site detection traditionally involve extensive unbiased molecular dynamics simulations, or more efficient but computationally still highly demanding biased MD alternatives that make use of enhanced sampling along low-frequency motions of the protein structure. Here we utilize a new variant of the recently introduced and orders of magnitude faster, MD-free COVIB algorithm to identify cryptic pockets efficiently in two challenging targets, the KRAS oncoprotein and the N-terminal domain of the STAT5B transcription factor. Using experimental and AI-modelled structures, respectively, we identify numerous cryptic sites, some of which have not yet been reported. The sites are prioritized and validated with our pharmacophore optimized photoaffinity-tagged fragment library via MS-based peptide mapping and NMR spectroscopy. Besides identifying new, targetable regions on the surface of these important oncotargets, we have established a protocol combining COVIB-based cryptic pocket detection and photoaffinity-based experimental peptide mapping as a robust approach for exploring unique and tractable cryptic pockets.

## Introduction

Traditionally, small-molecule drug discovery focuses on the orthosteric binding site, which is the specific pocket on a protein where endogenous ligands, substrates, or cofactors bind to initiate a biological response.^1^ While targeting these sites has yielded a vast amount of therapeutics, the approach is increasingly hindered by the fact that orthosteric sites are highly conserved across protein families, resulting in selectivity problems and the subsequent risk of off-target toxicity.^1–3^ Furthermore, orthosteric drugs must compete with the high-affinity natural ligand that is often present in the cell at high concentrations, like in the case of ATP in kinases.^4^

To circumvent these problems, orthosteric inhibitors must have extremely high affinity or have to be administered in high doses, however both options come with their own complications.^1,4^ High affinity is usually achieved by increasing lipophilicity, which compromises the pharmacokinetic profile and selectivity, as hydrophobic compounds tend to bind proteins nonspecifically and aggregate at high concentrations,^5,6^ while high doses can also lead to selectivity issues, as the drug can bind to structurally similar active sites, ultimately leading to toxicity issues and adverse drug reactions, which remains a primary reason for high attrition rates in clinical trials.^4,7^

The limitations associated with orthosteric sites have driven growing interest in alternative strategies, particularly the exploitation of allosteric sites located at static or cryptic pockets. Allosteric drugs bind to sites on the target that are topographically distinct from the orthosteric site, and work by inducing structural changes in the protein, thereby modulating its activity.^1,3,4^ Compared to orthosteric drugs, allosteric drugs can influence signaling less drastically, their effects being practically “capped”.^1,3,4,8^ As allosteric sites are rarely conserved across protein families, it is easier to achieve higher degrees of selectivity, reducing or eliminating the possibility of toxicity problems.^1,3,4,8^

Cryptic pockets are traditionally defined as binding pockets that are not visible in apo (unliganded) structures of the target protein.^9,10^ Although cryptic sites are frequently revealed in ligand-bound structures, the dynamic nature of proteins suggests a more accurate definition: they correspond to pockets that are minimally populated or exhibit low volumes within the conformational ensemble of the apo state.^9–11^ Ligands targeting cryptic sites can induce conformational changes, and thus influence the activity of the target.^12^ This suggests that identifying relevant cryptic sites can enable the targeting of proteins that are considered undruggable, thereby expanding the ‘druggable genome’.^9^ Several clinically approved drugs, such as KRAS^G12C^ inhibitors sotorasib^13^ and adagrasib^14^ which target the Switch-II cryptic pocket, or the BCR-ABL1 inhibitor asciminib^15^ targeting the myristoyl allosteric site, exemplify the successful targeting of allosteric and cryptic binding sites. Further examples include GPCR allosteric modulators like maraviroc^16^ or cinacalcet,^17^ as well as type III kinase inhibitors like trametinib or cobimetinib.^18,19^ There are also several drugs currently in clinical trials like the SHP2 allosteric inhibitor TNO155,^20^ or the KRAS^G12D^ inhibitor MRTX1133^21^ targeting the Switch-II cryptic pocket.

These successes highlight the therapeutic potential of targeting cryptic and allosteric sites and facilitate the need for robust computational and experimental methods capable of identifying these “hidden” pockets. In general, binding pocket detection can employ a range of software solutions, which usually follow two main schools of thought. Geometric or grid-based algorithms like Molecular Discovery’s BioGPS^22,23^ or Schrödinger’s SiteMap^24,25^ detect binding sites against regular three-dimensional grids, based on geometric features/groupings or energy terms calculated with simple (mostly monoatomic) probes. By contrast, probe-based approaches like FTMap employ more complex probes (usually small molecules with different characteristics, incl. common solvents) to search the protein surface for favorable binding locations with docking, Monte Carlo simulations or other methods.^26,27^ Our recent experiences in the fields of fragment library design,^28,29^ and the detection of allosteric GPCR binding sites^30,31^ have prompted us to rely on the FTMap approach in this work for binding site detection.

While the above algorithms provide a solution for detecting pockets on the surface of a static structure, the detection of deep cryptic pockets involves another, crucial step: efficient mapping of the conformational space of the flexible protein target. As an established, but compute-intensive methodology, molecular dynamics (MD) simulations were applied successfully to discover druggable cryptic pockets of several important therapeutic target proteins, leading to validated lead molecules against e.g. the CB1 cannabinoid receptor^32^ or the β₂-adrenoceptor.^33^ Besides the option of running extensive MD simulations, cryptic pocket detection usually relies on techniques that, e.g., enhance sampling along the low-frequency collective motions (normal modes) of the protein structure.^34^

The recently introduced COVIB (co-vibrational modes) algorithm utilizes superpositions of a selected subset of a large number (200-500) of low-frequency all-atom normal mode eigenvectors.^35^ COVIB originally provided a directed co-vibrational mode comprised of a particular superposition of 50-100 of the lowest normal modes that best approximates the displacement vector of two experimentally resolved superposed end-states.^35^ Here we introduce a new application of random co-vibrational modes utilizing random superpositions of 20-50 of the most *collectively flexible* low-frequency normal modes to shake up a single experimental structure to probe its conformational neighborhood.

Protein conformations explored by the random-COVIB workflow was next mapped by FTMap identifying available binding pockets. This protocol was applied against challenging drug targets exemplified by KRAS, a widely relevant oncological target and the N-terminal domain of the transcription factor STAT5B implicated in a wide range of cancers, most importantly leukemia.^36^ In both cases, we identify several cryptic pockets, some of which have already been successfully targeted during our recent efforts,^29^ as well as by others.^13,14,37^ Crucially, new sites were experimentally validated by our PhotoXplorer platform^28,29^ relying on the MS-based detection of photoaffinity-labeled small-molecule probes, and here we disclose the first reported binders of several yet unexplored cryptic sites on the surface of both targets.

## Methods

The schematic overview of the computational protocol is shown in Figure 1A.

**Figure 1.**
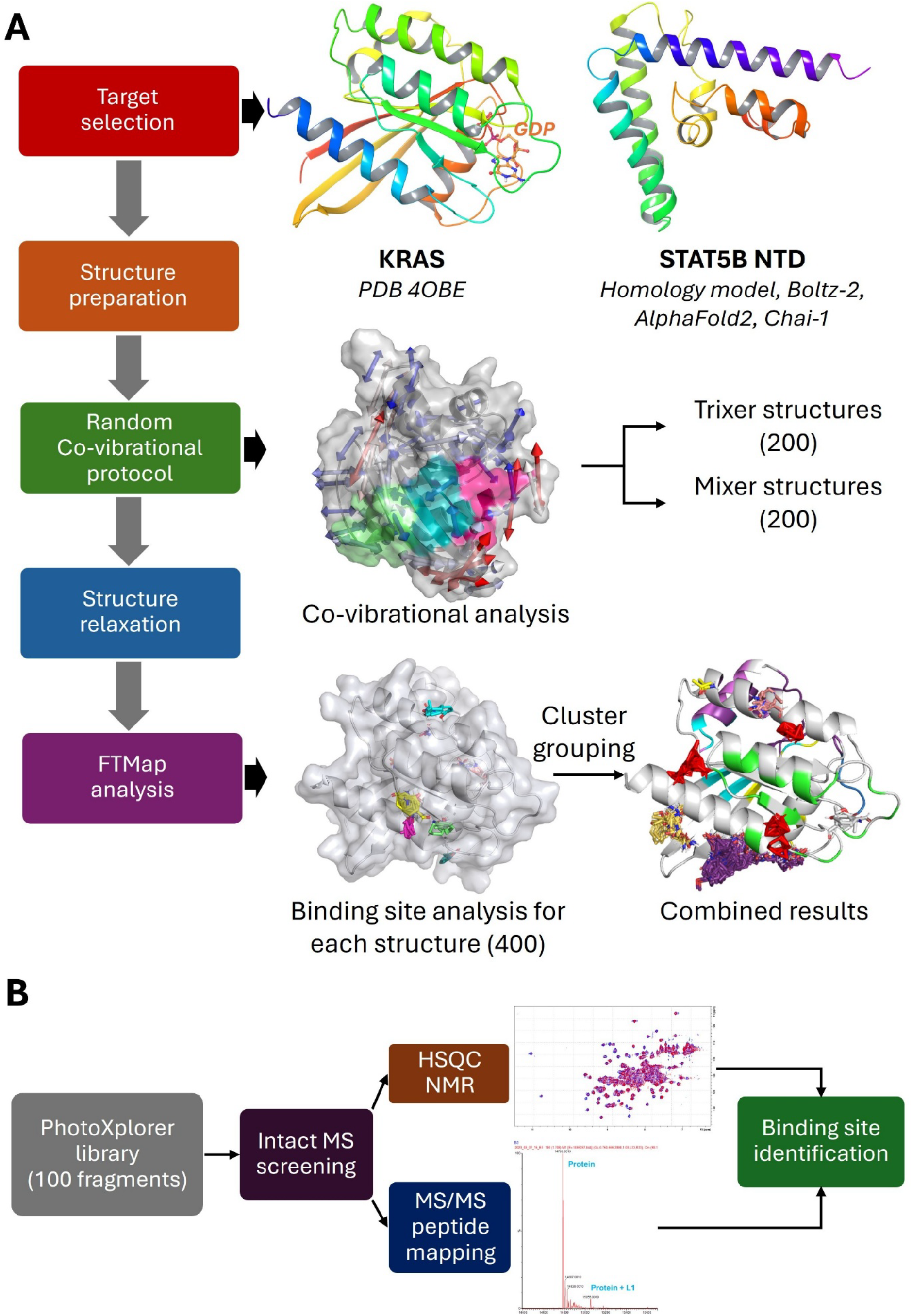
Overview of the full computational protocol (A) and experimental validation process (B).

### Input structures

For KRAS, the PDB structure 4OBE^38^ was prepared with Maestro’s Protein Preparation Workflow^39^ at a pH of 7.4 using default settings and deleting entries other than the protein.

For the STAT5B N-terminal domain, structures were generated by modeling, and included a legacy homology model^29^ (based on the STAT3 N-terminal domain structure 4ZIA^40^), Boltz^41^ (1 structure) and Chai^42^ (5 structures), while one structure was also acquired from the AlphaFold^43,44^ database (AF-P51692-F1 model). Protein homology modeling is the traditional approach which uses template-based alignment and physics optimization to generate structures based on an experimentally solved template.^45^ The AlphaFold database represents the first-generation deep learning revolution regarding structure prediction, as it processes evolutionary data through deep neural networks to iteratively model how physical parts of a molecule fold and interact.^43,44^ Boltz and Chai represent next-generation biomolecular foundation models that build upon the diffusion-based architecture of AlphaFold 3. While both utilize a denoising diffusion process to predict structures, they optimize for different pipelines: Boltz uniquely integrates a dedicated affinity module to predict binding strength at an accuracy approaching physics-based simulations, whereas Chai-1 leverages protein language model embeddings to reduce reliance on Multiple Sequence Alignments (MSAs), significantly accelerating inference times for single sequences.^41,42,44^

Structures were prepared with Maestro’s Protein Preparation Workflow at a pH of 7.4 using default settings. Residues 1-128 were used for further analysis (NTD): pairwise backbone atom RMSDs were calculated using Biopython^46^ for all structure (see Supplementary Table S1), and one structure from each source was selected, resulting in a total of four structures. In the case of the Chai structures, the one with the highest average RMSD value compared to the homology, AlphaFold and Boltz structures was selected.

### COVIB methods

For the reference conformation, the potential energy is expanded as

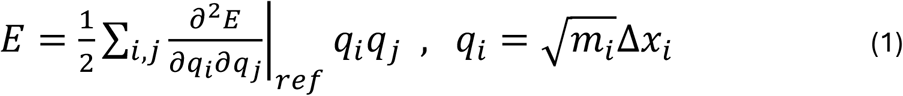

in mass-weighted Cartesian displacements 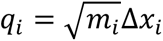, where m_ᵢ_ is the mass of the ⌊(*i* − 1)/3⌋*^t^*^ℎ^atom, *N* is the total atom count, and the mixed second partial derivatives form the elements **H**_ᵢⱼ_ of the mass-weighted Hessian. For a general-purpose force field (AMBER with a Generalized Born implicit solvent in this work), the reference structure is first energy-minimized. Because the full all-atom Hessian is too large to store and diagonalize, only the *M* lowest-frequency mass-weighted eigenvectors **v**_ⱼ_ are computed with an iterative Krylov-type solver,^35^ using purely implicit Hessian–vector products,

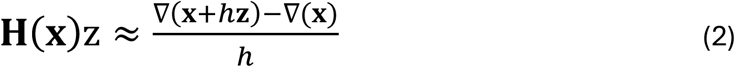

obtained by finite differencing of the analytic gradient. The resulting **v**_ⱼ_ are transformed back to non-mass-weighted Cartesian coordinates for all subsequent atomic-displacement operations. For both applications below we computed the first *M*=500 low-frequency all-atom modes.

#### Directed (allosteric) co-vibrational mode

When two experimental end-state structures are available, the all-atom displacement vector **f** from a chosen reference toward the other state (after optimal superposition) is expanded in the eigenvector basis of the reference,

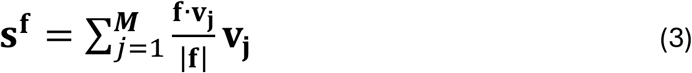

with weights equal to the direction cosines between f and each normalized eigenvector. Physically, **s^f^** is the instantaneous response of the reference harmonic oscillator when perturbed toward the other end-state. In our experience, COVIB modes with even a moderate *M* ≈ 50 can approximate **f** significantly better than any single normal mode.

#### Random co-vibrational modes for cryptic-pocket discovery

In our new random COVIB approach, the same construction can be used in the *absence* of a second experimental structure to efficiently explore the conformational landscape around one given structure. After computing the first *M* = 500 all-atom eigenvectors of the reference, we selected the subset S ⊆ {1, …, *M*} of the most collectively flexible modes — operationally, the low-frequency modes whose displacement is delocalized over a large fraction of the protein, as quantified by their collectivity index such as the so-called mode collectivity index,^47^ or a simple geometric mean to penalize outliers. We then replace the deterministic projection weights in Eq. (3) by random coefficients,

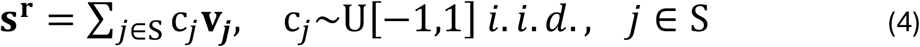

Each draw of {c_ⱼ_} yields a different random co-vibrational direction **s^r^**, along which the structure is then *flexed* using the curvilinear ZIG-ZAG protocol described below. Repeating this procedure with many independent random draws generates an ensemble of physically reasonable, force-field-relaxed conformations that collectively sample the soft degrees of freedom accessible to the protein. Applied to a panel of proteins, this random-COVIB protocol predicted a wide range of cryptic pockets, including pockets that had not been observed in any prior experimental structure; and the predicted pockets were subsequently confirmed experimentally, validating the use of random co-vibrational mixing as an efficient cryptic-pocket-discovery engine.

#### Curvilinear COVIB mode displacement and structure ffexing (ZIG-ZAG)

Because the mode vectors represent linear displacement along any co-vibrational direction **s^f^** or **s^r^**, the perturbation rapidly distorts hard internal degrees of freedom and introduces serious local geometric deformations and steric clashes. We therefore advance along the mode vector using the ZIG-ZAG algorithm,^35^ which iterates short Cartesian displacements along the mode (“zig”) followed by a few cycles of local energy minimization terminated when the gradient norm drops below 1 kcal mol^-1^ Å^-1^ (“zag”). The partial “zag” minimization relaxes bond-length and bond-angle strain and relieves clashes while allowing progress along the soft, collective degrees of freedom, yielding smooth, large-amplitude curvilinear trajectories.^35^ Random-COVIB employs the same ZIG-ZAG algorithm to probe the reference structure’s local conformational neighborhood efficiently. It should be noted that while the COVIB trajectories are curvilinear and can extend to substantial 2-5 Å amplitudes, the farther the structure is flexed from its starting conformation the higher its energy increases. Therefore, we always applied structure relaxation to present candidate conformations for evaluating the existence of cryptic pockets. We applied two different versions of random-COVIB, one called trixer modes where the random coefficients for mixing could only have 3 different values (-1, 0, 1) and mixer modes where the coefficients could take any value in the [-1,1] interval.

#### COVIB computational protocol

Our code to compute co-vibrational modes was originally added in-house to the low-mode conformational search module (LMOD)^48^ in the AmberTools24 release of the Amber molecular dynamics program package,^49–51^ but now it is publicly available in AmberTools25 and more recent releases. We ran our calculations on a Linux workstation, utilizing 32 4-GHz AMD Ryzen CPU cores, using the ff14SB^52^ force field, and a generalized Born implicit solvation model^53^ with infinite cutoffs. Data files and scripts with instructions to reproduce our results are publicly available.^54^

### Binding site mapping

Two representative frames were selected from each of 100 trixer and 100 mixer modes, resulting in a total of 400 frames per protein structure. The selected frames for each mode were the highest amplitude endpoints of the mode “swing” in both directions. Each frame was relaxed using the FastRelax^55^ protocol in PyRosetta^56^ with a constraint, so that the conformation of the protein backbone may not differ from the input structure, only the sidechains could move. Atlas, the command line implementation of the FTMap algorithm^57^ was then used to identify possible binding sites on the relaxed structures. For reference, detection of possible binding sites was also carried out for the unmodified, prepared (experimental or modeled) input structures (labeled later as unrefined structures).

To note, FTMap identifies binding sites based on the energetically favorable locations of probe clusters. In many cases, the same location is identified across a range of protein structures, resulting in dense probe clusters. For a given protein, to filter out redundancies, the probe clusters identified on co-vibrated structures (labeled as cryptic clusters, **C**) were ignored if the center of mass of the cluster was within 10 Å of the center of any identified cluster on the appropriate reference structure (labeled as open clusters, **O**). The remaining COVIB clusters were grouped using density-based clustering found in Scikit-learn^58^ (epsilon = 2 Å). Visualization has been done using PyMOL.^59^

### Experimental validation

A 100-member library of pharmacophore optimized fragments tagged with the diazirine photoaffinity handle was screened by mass spectrometry to prove binding to the corresponding targets by anchoring covalently upon irradiation (Figure 1B). The expression and purification of KRAS^G12D^ and STAT5B-NTD was reported previously.^29^ In case of KRAS^G12D^, the binding sites of the proven binders were then mapped by HSQC NMR to identify the amino acid residues interacting with the small molecule hits. Clustering these amino acids led us to the identification of binding pockets that could be compared to the computationally determined ones. In particular cases, binding sites were further confirmed by digestion experiments for both KRAS^G12D^ and STAT5B-NTD. The library design, synthesis and compound characterization was disclosed previously.^29^ The experimental results (MS, NMR) used for the validation of binding sites identified by the COVIB-FTMap protocol are detailed in the Supplementary Information of this paper (Pages S6-S33)

#### Sample preparation for STAT5B-NTD illumination

STAT5B protein in 7 µM concentration in pH 7.4 PBS buffer was pre-incubated with 0.17 µL PhP (in 100 mM DMSO solution) for 1 h at room temperature in dark. Irradiation was carried out at 4 °C for 10 min using 365 nm UV lamp.

#### Intact mass spectrometry (MS) screening

The PhotoXplorer library^29^ was screened against KRAS^G12D^ as follows. Using a Labcyte Echo^®^ 555 Liquid Handler, 150 nL of fragment solution (20 mM) was transferred into a Greiner 384 low-volume plate (white) to prepare the probe plate. The probe plate was placed on ice. 15 μL of protein stock solution (1 μM KRAS^G12D^ (GSK/GenScript) in PBS buffer) was dispensed into wells containing fragments. The plate was left on ice for 15 min for incubation, then irradiated at 302 nm for 10 min, and centrifuged (1000 rpm, 1 min) to remove any bubbles. The plate was then analyzed by LCMS-TOF mass spectrometry (Agilent 1200 series liquid chromatography with Agilent Bio-HPLC PLRP-S (1000 Å, 5 µm × 50 mm × 1.0 mm, PL1312-1502) reverse phase HPLC column at 70 °C equipped with an Agilent G6224 time-of-flight (ToF), see Supplementary Information for the full protocols). The deconvoluted spectra were analyzed using R Studio software. Spectra of the hits are reported in the Supplementary Information.

For screening against STAT5B-NTD, we have used an UHPLC-MS system that consisted of a Waters ACQUITY UPLC I-Class setup coupled with a Waters ACQUITY UPLC Peptide BEH C18 Column (130 Å, 1.7 µm, 2.1 mm × 100 mm), connected to a Waters Xevo G2-XS QT-ToF instrument equipped with a Waters Z-spray ESI source. During the analysis, the column temperature was maintained at a constant 60 °C, and a sample volume of 3 µL was injected for each analysis. Data acquisition was conducted in positive ion mode within the *m/z* 100–2000 (mass-to-charge ratio) range. For the full protocol and spectra of the hits, see the Supplementary Information.

#### Binding site identification by LC-MS/MS peptide mapping — STAT5B-NTD

In follow-up of the intact MS screening, the fragment hits were further analyzed by a Triple TOF 5600+ hybrid Quadrupole-TOF LC/MS/MS system, after digesting the resulting fragment-protein complexes by Trypsin/Lys C mix. Data acquisition and processing were performed using Analyst TF software version 1.7.1 (AB Sciex Instruments, CA, USA). Chromatographic separation was achieved on the Discovery^®^ BIO Wide Pore C-18-5 (250 mm × 2.1 mm, 5 μm, 300 Å) HPLC column. MS/MS spectra were obtained on the 8 most abundant parent ions present in the TOF survey scan with the Information Dependent Acquisition (IDA) mode, and peaks were evaluated with PeakView^®^ (version 2.2, Sciex) and Biologics Explorer (version 3.0.3, Sciex). Sample preparation and data acquisition protocols, as well as the spectra of the reported hits, are available in the Supplementary Information.

#### Binding site identification of KRAS^G12D^ hits by LC-MS/MS peptide mapping

Modification sites were determined by proteolysis and reversed-phase LC-MS peptide mapping. Briefly, proteins were enzymatically digested in 25 mM NH_4_HCO_3_ solution after buffer exchange using Amicon Ultra-0.5 mL Centrifugal Filter units (10 kDa, Merck Millipore). Trypsin-LysC mixture and ProAlanase (Promega Corporation, Madison, USA) were used for enzymatic digestion. Protein samples were reduced by dithiothreitol at 37 °C for 30 min. After reduction, proteins were digested using 1:50 enzyme:protein ratio or 4 h at 37 °C, followed by an additional short incubation with dithiothreitol (5 min, 37 °C). Overnight digestion was performed by Trypsin-LysC mixture in 50 mM NH_4_HCO_3_ solution at 37 °C. Tryptic digestion was stopped by adding formic acid in a final concentration of 0.2% (V/V). ProAlanase digestion was performed for 4 h at 37 °C in 50 mM HCl and was stopped by heating at 90 °C for 10 min. After digestions an additional short incubation with dithiothreitol was repeated (5 min, 37 °C).

Mass spectrometric experiments were performed on a high-resolution hybrid quadrupole-time-of-flight mass spectrometer equipped with a cyclic ion mobility separator (Waters Select Series Cyclic IMS, Waters Corp., Wilmslow, U.K.). Chromatographic separation was performed using a Waters Acquity I-Class UPLC system, coupled directly to the mass spectrometer. Waters Acquity CSH Peptide C18 UPLC column (2.1 × 150 mm, 1.7 µm) was used for chromatography. Gradient elution was performed under the following parameters: eluent A: 0.1% formic acid in water, eluent B: 0.1% formic acid in acetonitrile; flow rate: 300 µL/min; column temperature: 60 °C; gradient: 2 min: 2%B, 80 min: 45%B, 81 min: 85%B. HDMS^E^ experiments were performed using a single-pass cyclic ion mobility separation and fragmentation in the transfer cell with collision voltage ramping. MS data acquisition was performed with the following parameters: *m/z* 50–2000, V-mode, scan time: 0.3 s, single Lock Mass: leucine enkephalin; low energy: 6 V, high energy: ramping 19-45 V. BiopharmaLynx 1.3.5 software (Waters Corp., Wilmslow, U.K.) was used to for data analysis. Spectra of the hits are reported in the Supplementary Information.

#### Binding site identification of KRAS^G12D^ hits by NMR spectroscopy

NMR samples contained 70 μM ^15^N-labeled GDP-bound KRAS^G12D^ protein, 100–600 μM binding partner, 10 mM MgCl_2_, 3 mM NaN_3_, in PBS buffer, 5% DMSO*-d_6_*, 7% D_2_O, 0.5% DSS and the pH was set to 7.4. ^1^H,^15^N-SOFAST-HMQC (fast version of ^1^H,^15^N-HSQC) NMR spectra were acquired at 298 K on a Bruker AVANCE III spectrometer (Bruker Biospin, Rheinstetten, Germany) operating at 700.05 MHz for ^1^H and 70.94 MHz for ^15^N, equipped with a 5 mm Prodigy TCI H&F-C/N-D, z-gradient probe head. Temperature was calibrated by standard methanol solution. The chemical shifts were referenced with respect to the ^1^H-resonance of an internal DSS standard, while ^15^N chemical shifts were referenced indirectly via the gyromagnetic ratios according to the IUPAC conventions. All NMR data were processed with Bruker TOPSPIN 3.6 and analyzed in POKY software.^60^ The shifted crosspeaks were compared to the free KRAS^G12D^ chemical shifts.^61^ Spectra of the hits are reported in the Supplementary Information.

#### Dual clustering protocol for the binding site identification of PhotoXplorer hits from CSP data

PhotoXplorer binding sites were identified by the previously developed PhotoFold protocol.^62^ In short, ^15^N HSQC NMR spectra were recorded for samples containing 70 μM ^15^N-labeled GDP-bound KRAS^G12D^ and 100-600 μM binding partner. Chemical shift perturbation (CSP) values were determined for 13 different PhotoXplorer hits by comparing the amide peaks with the control K-Ras spectrum. For the clustering protocol, we used the crystal structure of the GDP-bound form of WT KRAS (4OBE).^38^ Binding sites were identified using a two-step clustering protocol that combined CSP data with the 3D structure of the protein. First the center of binding sites was determined using average-linkage hierarchical clustering based on the distance of the center-of-mass (COM) coordinates of all residues with CSP above a value defined by a parameter (*s*). The binding criterion is defined as the distance *D*_0_. The collected centers of each binding site are again clustered using the K-means method.^63^ The separation of clusters is measured by the Silhouette score.^64^ Parameters *s* and *D*_0_ and the final number of clusters are determined by an optimization procedure.

## Results

### Computational protocol

We started the process by selecting the KRAS structure 4OBE and four modeled structures of the STAT5B N-terminal domain: a legacy homology model based on the STAT3 N-terminal domain (PDB: 4ZIA^40^), and further ones acquired from the AlphaFold database (AF-P51692-F1 model) and modeled with Boltz-2 and Chai-1. After preparing the structures, the random COVIB protocol was applied to each protein. We computed the first 500 low-frequency all-atom normal modes and then selected the 20 collectively most flexible modes as the basis vectors for COVIB mode generation. See the Methods section for details. 100 mixer and 100 trixer co-vibrational modes were generated per structure, out of which the extrema were extracted from each mode, resulting in a total of 400 structures ((100+100)×2) in the case of KRAS and a total of 1,600 structures in the case of the STAT5B N-terminal domain (4×((100+100)×2)). Each structure was then relaxed before FTMap analysis. It is worth mentioning one of the advantages of the method here, which is that it is significantly simpler and faster compared to MD simulations.

The FTMap calculations were run on all 2,000 structures resulted from the COVIB protocol alongside the five input structures for reference. Each calculation resulted in probe molecule clusters mapped on the protein surface, locating possible binding sites. For a given input structure, these results were then combined, and the clusters were grouped based on proximity from each other. Two types of groups were distinguished: probe clusters identified on the reference structures (also labeled as **O** for open clusters), and probe clusters identified on the structures resulted from the COVIB protocol which were not near (within 10 Å) the center of any of the open clusters (also labeled as **C** for cryptic clusters). This way, cryptic clusters map possible binding sites which are not identifiable by analyzing only the unmodified X-ray structure.

### Mapping of protein-protein binding sites of KRAS

To evaluate the performance of the COVIB protocol, we have computed 400 structures corresponding to the extrema of 100 mixer and 100 trixer co-vibrational modes of the KRAS oncoprotein (PDB: 4OBE^38^), an important driver of several oncological indications, with a variety of well-studied protein-protein binding sites.^29^ We have mapped the surfaces of each structure with Atlas, a command line implementation of the popular FTMap binding site detection algorithm (resulting probe clusters are labelled with **C** for cryptic clusters), along with the surface of the input structure 4OBE (labelled with **O** for open clusters). The locations of the probe clusters are shown in Figure 2A-D, while the corresponding experimental hits are shown in Figure 2E. The number of probe molecules, as well as the corresponding binding sites are summarized in Supplementary Table S2.

**Figure 2.**
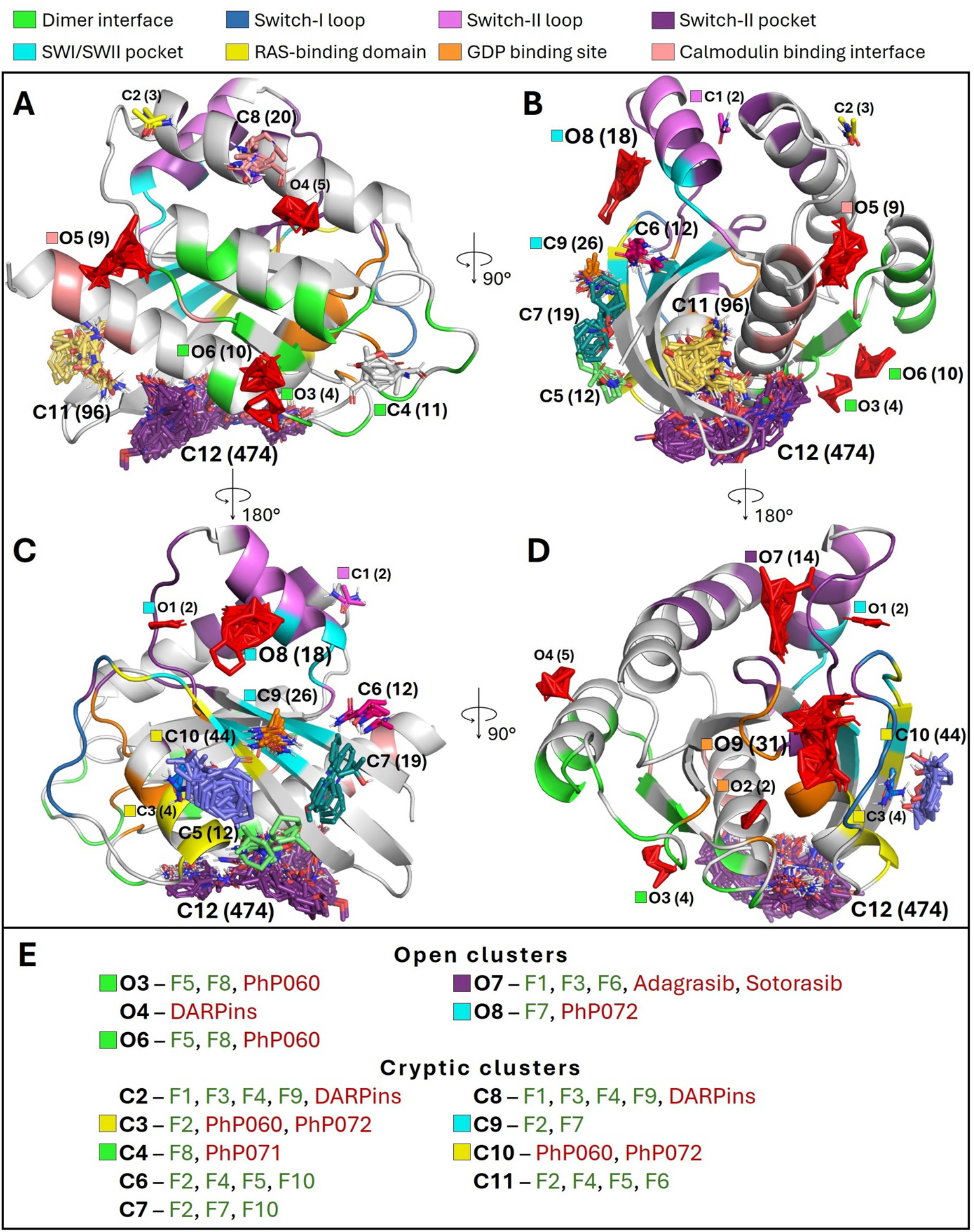
(A-D) KRAS protein structure (PDB ID: 4OBE) with the locations of the probe molecule clusters identified by Atlas in four different viewing angles (probe clusters obstructed by the protein are omitted). The residues on the protein structure are colored based on known binding sites, as explained in the top legend. Open clusters are uniformly colored red, while cryptic clusters are colored by size (following the default palette of PyMOL). Each cluster is labeled with its corresponding ID (cf. Table S2), and the number of probe molecules being included in brackets. A colored square is located next to the label, if the cluster corresponds to a known binding site. with the coloring based on the known site as explained in the top legend. The GDP molecule from the input structure is omitted for clarity, but the residues locating the GDP binding site is colored in orange. (E) Experimentally confirmed site(s)/hit(s) from earlier works (red) and this work (green),identified by the appropriate clusters.

Calculations on the reference protein structure resulted in nine probe clusters (Figure 2A-D, labeled O1-O9), from which seven located known binding sites or interfaces (O1, O2, O3, O6, O7, O8, O9). The binding site of the native ligand GDP is also identified with clusters O2 and O9, the latter of which contains the greatest number of probe molecules out of all unrefined clusters, nicely showcasing the primary ligand-binding hot spot. This trend also continues with the open clusters containing the second and third most probe molecules (O8 and O7), which locate the SWI/SWII pocket and Switch-II pockets, respectively, the main druggable sites exploited by most of the known KRAS inhibitors. Open clusters O4 and O5 were located on the protein surface not corresponding to any well-known binding sites (although O5 locates a less well-known site that was described to bind calmodulin).^65,66^ None of the open clusters located the RAS-binding domain, which could be explained by it being a protein-protein interface with a flat surface, making it difficult to map with the default settings of Atlas. We note here that as the unrefined structure was used for visualization, uncertainties may be present in locating binding sites, as the structural shift between the input and co-vibrated structures can result in slight shifts in β-sheet spacing.^67,68^

Binding site prediction was also done for the 400 structures resulting from the co-vibration analysis. As the protein structure is slightly modified during the calculations, we expected shallow and cryptic sites to be also identified by probe molecules. After combining the resulting clusters, a total of twelve unique cryptic clusters were identified (Figure 2A-D, labeled C1-C12). Out of the twelve cryptic clusters, five located known binding sites or interfaces (C1, C3, C4, C9, C10).

Clusters C2 and C8 are both located at helix α3 next to the Switch-II pocket, with cluster C2 being at the C-terminal end. Clusters C5, C6 and C7 are located next to each other, with cluster C5 between the C-terminal end of helix α1 and sheet β2, next to the RAS-binding domain, cluster C6 at sheet β1, next to the SWI/SWII pocket, and cluster C7 between sheets β1, β2 and β3, also next to the SWI/SWII pocket. Clusters C11 and C12 are the two largest cryptic clusters, with 96 and 474 probe molecules respectively, and neither locates well-known binding sites. Cluster C11 is located between helix α5 and sheets β2 and β3, while C12 is located between helixes α1 and α5 and sheet β2.

Next, we compared the results with other computational studies for binding site prediction. Gomez-Gutierrez et al. performed accelerated molecular dynamics calculations, and identified eight prospective allosteric sites, which were labeled as sites P1-P8.^69^ We were able to correlate a cluster to each of the prospective allosteric sites identified in that work, e.g. clusters O5 (site P3), C8 (site P8), C11 (site P6) and C12 (site P5), which have not located any well-known binding sites or interfaces. Specifically, sites P5 and P6 locate the two most populous clusters C12 and C11, respectively.^69^ Site P5 is described as an undruggable site, while the druggability of site P6 is labeled as difficult. They also mention that binders of site P5 (which is the part of the recognition interface of the cysteine-rich domain of the Raf protein) can act as PPI inhibitors of KRas/Raf.^69,70^ Sites P1, P2, P4, P7 located the same sites as clusters O8, O7, C3+C10 and O6, respectively. Vithani et al. performed weighted ensemble molecular dynamics simulations with inherent normal modes and all-atomic simulations with and without several cosolvents, including xenon.^34^ The authors have identified eight high-affinity xenon-binding clusters: here, we labeled the xenon-binding clusters as Xe1-Xe8, with Xe1-Xe4 corresponding to the original labeling, Xe5 being the cluster between the N-terminal end of helix α1 and C-terminal end of sheet β4, Xe6 being the cluster at the N-terminal end of helix α4, Xe7 being the cluster between helix α4 and the N-terminal end of sheet β6 and lastly, Xe8 being the cluster at the N-terminal end of helix α5 (these were not annotated in the original work). Out of the eight xenon-binding clusters, we were able to correlate seven with identified clusters, with the exception being Xe5. Xenon-binding clusters Xe1, Xe2, Xe3, Xe4, Xe6, Xe7 and Xe8 located the same sites as clusters O7, O1+O8, C2, C1, C4, O6 and C12, respectively.^34^ Kapoor et al. used extensive molecular dynamics simulations on HRAS, KRAS and NRAS, where they identified pocket p5 (nb. not the same as P5 earlier) among others as a particularly interesting site, which locates the same site as the most populous cluster C12 here.^71^ Alongside site p5, sites p1, p2, p3 and p4 located the same as clusters O1+O8, O7, O5+C2+C8 and C10, respectively.^71^

The identified sites from computational studies corresponding to the open and cryptic clusters are summarized in Supplementary Table S2.

We have also compared experimentally validated results with our identified probe clusters (Figure 2E). The two FDA-approved KRAS inhibitors, sotorasib^13^ and adagrasib^14^ both bind to the Switch-II pocket, corresponding to cluster O7. Our recent work about mapping protein binding sites by photoreactive pharmacophores also included three experimentally validated (by tryptic digestion followed by MS/MS and by HSQC NMR, spectra are shown in the SI) hits against KRAS^G12D^.^29^ Fragment PhP060 labeled two sites, the RAS-binding domain, corresponding to clusters C3 and C10, and the dimer interface, with its position corresponding to clusters C3 and C6. Fragment PhP071 labeled only the dimer interface (C4), while fragment PhP072 labeled two sites, the Switch-II pocket (C7) and the RAS-binding domain (C3 and C10). Bery et al. reported two designed ankyrin repeat proteins (DARPins) that can inhibit KRAS by binding to the allosteric lobe that includes residue H95.^37^ The binding interface of the DARPins K13 and K19 corresponds to clusters O4, C2 and C8. Buhrman et al. performed a combination of multiple solvent crystal structures (MSCS) and computational solvent mapping (FTMap) to determine binding site hot spots in HRAS, KRAS and NRAS.^72^ The MSCS method resulted in eight clusters. Clusters 1, 2, 3, 4, 6, 7 located the same sites as our clusters O7, C8, C3, O2, O9 and C12, respectively. Cluster 8 also located the same site as cluster O9, while cluster 5 has no corresponding identified cluster from this work. The FTMap results correlated with our results with the reference structure (corresponding clusters are clusters O3, O4, O5, O6, O7, O8).^72^

To expand the number of experimentally identified binding sites or interfaces, we have conducted further experiments. HSQC NMR screening of our photoaffinity-tagged pharmacophore fragment library^29^ resulted in 10 hits, labeled as F1-F10 (spectra are shown in the SI). The binding sites of the hits were explored identifying their labeling site and their contacting residues by HSQC NMR. Computational binding site analyses were constrained by these experimental data and revealed the same sites as 12 clusters (O3, O6, O7, O8, C2, C3, C4, C6, C7, C8, C9, C11), out of which clusters C6 and C7 were not located by other methods before. Interestingly, no computational study or experimental work have identified the same site as cluster C5. For all clusters, the corresponding known site and identified sites from computational and experimental studies are summarized in Supplementary Table S2.

### Discovering cryptic sites on the STAT5B N-terminal domain

After evaluating the performance of the protocol on KRAS, we were encouraged by the results to adapt the calculations to a novel target, the N-terminal domain (NTD) of the transcription factor (and prominent leukemia driver) STAT5B. As there is much scarcer information on this target than KRAS (the only two well-established binding sites being the handshake dimer interface and the Ni^2+^ – mediated interface),^40,73^ the focus was mainly to identify possible novel targetable cryptic sites in a prospective manner.

Four structures were used as input, one legacy homology model based on the STAT3 N-terminal domain (PDB: 4ZIA^40^), one acquired from the AlphaFold database, one resulted from the Boltz^41^ algorithm, and one resulted from the Chai^42^ algorithm. Pairwise backbone atom RMSD values are below 5 Å for any pair among these structures (see Supplementary Table S1), showing that the overall global fold is conserved across all methods. Yet, the four methods introduce different biases into their generated structures: the homology model is heavily biased toward the reference template state, while AlphaFold is biased toward the dominant conformation in its PDB training data. In the meantime, both Chai and Boltz utilize diffusion-based sampling that inherently generates structural variance: Chai’s variance often reflects prediction uncertainty for novel sequences, whereas Boltz is trained to capture dynamic ensembles and can consequently oversample local conformational flexibility in isolated apo predictions. Consequently, while the global fold is conserved, there are significant structural deviations at the local loop and side-chain levels. For example, the local RMSDs for residues 51-98 (two long, parallel helices) diverge by up to 3.2 Å between the homology and Boltz models. Ultimately, these side-chain rearrangements alter the surface topology, leading to the identification of different potential binding sites and probe clusters depending on the source structure. By utilizing an ensemble of these four structures, we can distinguish template-induced biases from physically realistic conformational states, providing a more comprehensive map of the potential binding sites. The locations of the open clusters and cryptic clusters on all four structures are shown in Figure 3.

**Figure 3.**
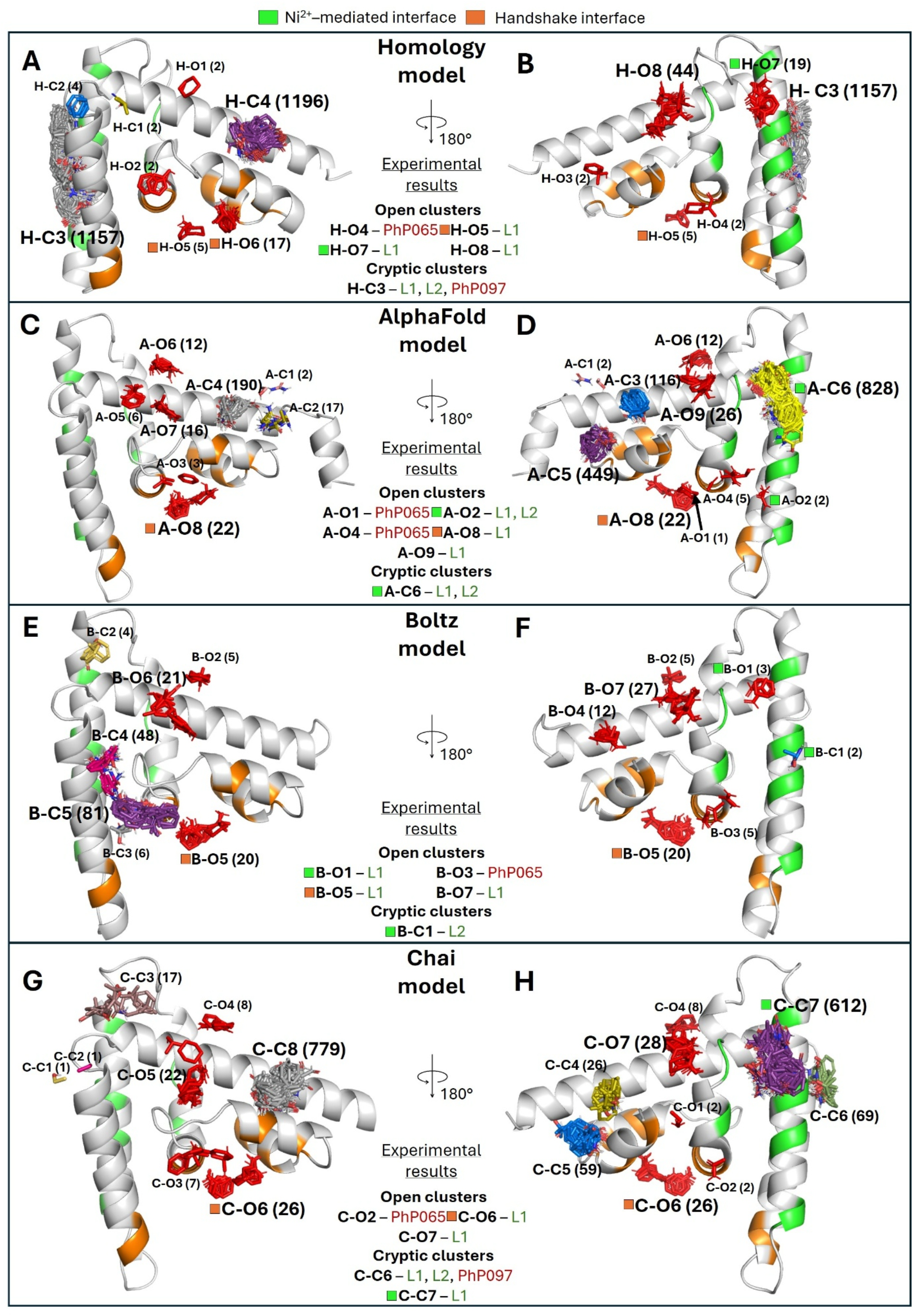
STAT5B N-terminal domain structures from all four sources: (A-B) Homology model, (C-D) AlphaFold model, (E-F) Boltz model, (G-H) Chai model, with the locations of the probe molecule clusters identified by Atlas in two different viewing angles (probe clusters obstructed by the protein are omitted). The residues on the protein structures are colored based on known binding site information, with green and orange corresponding to the Ni^2+^ – mediated interface and the handshake dimer interface, respectively. Open clusters are uniformly colored red, while cryptic clusters are colored by size (following the default palette of PyMOL). Each cluster is labeled with its corresponding ID (cf. Table S3), and the number of probe molecules being included in brackets. A colored square is located next to the label, if the cluster corresponds to a known binding site. with the coloring based on the known site as explained in the top legend. In the middle, experimentally identified hit(s) from earlier works (red) and this work (green) that bind to the respective sites are shown.

Calculations on the reference structures resulted in eight clusters on the homology structure (Figure 3A-B, labeled from H-O1 to H-O8), nine on the AlphaFold structure (Figure 3C-D, labeled from A-O1 to A-O9), seven on the Boltz structure (Figure 3E-F, labeled from B-O1 to B-O7) and seven on the Chai structure (Figure 3G-H, labeled from C-O1 to C-O7). The handshake interface was located by H-O5, H-O6, A-O8, B-O5 and C-O6, while the Ni^2+^ – mediated interface was located by H-O7, A-O2 and B-O1, with the rest of the open clusters located elsewhere.

Binding site prediction on the 400-400 structures per source from the co-vibration analysis resulted in four cryptic cluster groups on the homology structure (labeled from H-C1 to H-C4), six on the AlphaFold structure (labeled from A-C1 to A-C6), five on the Boltz structure (labeled from B-C1 to B-C5) and eight on the Chai structure (labeled from C-C1 to C-C8). Regarding known interfaces, the handshake interface was not assigned to any cryptic clusters (as it was already covered by open clusters), while the Ni^2+^ – mediated interface was located by clusters A-C6 and C-C7 at the same location as H-O7 and B-O1, and by cluster B-C1.

To the best of our knowledge, no computational studies were conducted specifically for binding site detection against STAT5B-NTD, so we analyzed experimental hits against the target. Our earlier work about mapping protein binding sites by photoreactive pharmacophores included two experimentally validated hits against STAT5B-NTD: PhP065 and PhP097.^29^ PhP065 binds close to the cavity formed in the handshake dimer interface, located by clusters H-O4, A-O1, A-O4, B-O3 and C-O2, while PhP097 binds at the tip of the helices α6 and α7 next to the Ni^2+^– mediated interface, located by clusters H-C3 and C-C6.

MS-based screening of our photoaffinity labelled pharmacophore fragment library identified two hits: L1 and L2. Constrained computational analysis of their binding sites revealed that L1 labeled STAT5B-NTD at five different locations: at the handshake interface (H-O5, A-O8, B-O5 and C-O6), at the Ni^2+^-mediated interface (H-O7, A-O2, A-C6), at a location characterized by clusters H-O8, A-O9, B-O7 and C-O7, and lastly at the same location as PhP097 (H-C3 and C-C6). L2 labeled STAT5B-NTD at three different locations: two alternative sublocations at the Ni^2+^-mediated interface (A-O2, L1 vs. A-C6, B-C1), and at clusters H-C3 and C-C6 where PhP097 and L1 also bind.

Unsurprisingly for STAT5B-NTD, due to the lack of data, there are numerous clusters not corresponding to known sites or experimental hits (Figure 3). For all clusters on each four structures, the number of probe molecules are summarized in Supplementary Table S3, while corresponding clusters from other structures, known sites and identified sites from experimental studies are summarized in Supplementary Table S4.

## Discussion

Analysis on both targets has shown that our approach can identify numerous potential binding sites without the need for significant computational resources. On the extensively researched target KRAS, we have identified twelve possible binding spots not found on the reference structure, from which three clusters (C5, C6 and C7) were not identified before by computational studies or experimental results or were not corresponding to well-known binding sites or interfaces. Retrospective analysis revealed that our results align with previously identified binding sites from both computational and experimental studies, with all but one site corresponding to a cluster in our dataset. Crucially, clusters C6 and C7 correspond to locations where ligand binding has been experimentally validated here, suggesting that these may represent novel cryptic sites for targeting KRAS.

A conceptually related cryptic-pocket discovery strategy using normal modes was recently reported for KRAS^G12D^ using weighted-ensemble molecular dynamics with anisotropic-network-model (ANM) progress coordinates,^34^ and it is instructive to highlight several methodological differences with random-COVIB:

*(i) 1-3 modes vs. a random mixture of 20-50 modes.* The WE-MD/ANM approach is built on the coarse-grained ANM of the Cα trace and only the top ∼20 slowest modes are inspected. One to three of the most collectively flexible modes are then chosen to drive a simulation. Random-COVIB operates instead on the *all-atom* Hessian, computes several hundred (200-500) low-frequency eigenvectors, and then forms an ensemble of *random linear combinations* of the 20-50 most collectively flexible modes (Eq. (4)). Therefore, a single COVIB perturbation inherently samples a high-dimensional subspace of all-atom soft motion rather than severely limiting the search by including only 1-3 Cα ANM modes.
*(ii) MD-driven curvilinearization vs. ZIG-ZAG.* Because a normal-mode vector is a *linear* Cartesian atomic displacement, both methods face the problem of converting it into physically realistic, curvilinear, large-amplitude motion. WE-MD/ANM solves this by running explicit-solvent molecular dynamics simulations under the weighted ensemble path-sampling protocol, using the projection of the instantaneous simulation state onto the selected ANM modes as the progress coordinate and, therefore, curvilinearity is an emergent property of using MD, which is physically meaningful but the cost is substantial: the published pipeline consumes >400 µs of aggregate WE MD and ∼47 Orion cloud-days *for a single protein*, and ultimately requires xenon, ethanol, benzene or MRTX1133 cosolvents plus exposon, dynamic-probe-binding and probe-map post-processing to detect pockets retrospectively. Random-COVIB achieves curvilinear, force-field-relaxed motion *without MD* by means of the ZIG-ZAG algorithm, in which 10–20 alternations of an LMOD step along **s^r^** and a short minimization to gradient norm < 1 kcal mol⁻¹ Å⁻¹ relieve hard-coordinate strain and steric clashes while leaving the soft, collective motion essentially intact.^35^
*(iii) Role of the mode in driving motion.* In the WE/ANM pipeline the selected ANM modes are only progress coordinates; the actual displacements are produced by stochastic MD. In random-COVIB the random co-vibrational vector is *itself* the displacement direction along which the structure is flexed, making the procedure deterministic at each step and orders of magnitude cheaper. Finally, the WE/ANM study to date has demonstrated *retrospective* recovery of known KRAS pockets, whereas random-COVIB has been used *prospectively* to identify previously unknown cryptic pockets across multiple targets, several of which were subsequently confirmed experimentally.
*(iv) Computational complexity and efficiency* COVIB is significantly simpler and orders of magnitude faster than the WE-MD/ANM approach; the entire COVIB calculation of 100 trixer modes and 100 mixer modes of KRAS^G12D^ based on 500 low-frequency, all-atom normal modes took a total of ∼2 hours on 32 CPU cores, running on a desktop workstation vs. ∼47 Orion cloud-days.

For STAT5B-NTD, we have identified numerous new potential binding sites, with the site located by clusters H-C3 and C-C6 being a promising novel cryptic site, as experimental results have shown three ligands targeting it and cluster H-C3 being the overall second largest cluster with 1157 probe molecules.

The results demonstrate the effectiveness of our method in comprehensively mapping potential cryptic sites, facilitating the identification of promising candidates and targetable sites for computational drug discovery.

## Acknowledgement

The authors thank Elvin D. de Araujo and Qirat F. Ashraf (University of Toronto Mississauga) for their kind contribution of the STAT5B N-terminal domain. MS measurements by Aaron Keeley and László Petri are gratefully acknowledged. We thank Sándor Vajda (Boston University) and Dmitry Beglov (Acpharis Inc.) for granting access to the Atlas software package. This work was supported by the National Research Development and Innovation Office of Hungary [contracts FK146063 to D.B., NKKP 153477 and 152137 to G.M.K.], the Hungarian Academy of Sciences [grant no. LENDULET_2026-55] and the Schweizerische Nationalfonds (SNF) [MAPS 230109 to G.M.K.]. The work of D.B. and P. Á.-B. was supported by the János Bolyai Research Scholarship of the Hungarian Academy of Sciences. We acknowledge the HPC time and support of the Governmental Information Technology Development Agency, Hungary.

## Data and Software Availability

Protein structure and sequence data were obtained from public data depositories (Protein Data Bank for the KRAS structure with the accession code 4OBE, AlphaFold database for the STAT5B NTD structure with the ID of AF-P51692-F1, UniProt for the residue sequence of the full-length STAT5B protein). The structures of photoaffinity-tagged fragments are available in ref.^29^. Results obtained from the COVIB protocol (all runs with code, trajectories, PyMol sessions and video files) are available via Zenodo at https://doi.org/10.5281/zenodo.21266702.^54^ The COVIB methodology is also available in the AmberTools package (https://ambermd.org/AmberTools.php). Python scripts used in the evaluations are available via GitHub at https://github.com/keserulab. Visualization and the evaluation of the results were done using PyMol v2.5.0 (https://www.pymol.org/). Protein mapping was performed with Atlas, the commercially licensed command line implementation of FTMap through the kind contribution of Sándor Vajda and Dmitry Beglov (a free online version is available at https://ftmap.bu.edu/ for academic use).

## Conflict of interest statement

The authors declare no competing financial interests.

## Author Contributions

I.K. has developed the COVIB methodology and performed the COVIB computations. T.V.S. has performed the Atlas computations, mapping of binding sites and visualization. P.Á-B. and Z.O have performed experimental work. M.G. measured the NMR spectra, T.I., E.K.G. and J.T.B. performed MS measurements. D.B., I.K. and G.M.K. have conceptualized and directed the study. G.M.K. have supervised the project and provided funding. All authors have participated in writing the manuscript.

## Supplementary Information

Pairwise backbone atom RMSD values calculated between different STAT5B N-terminal domain structures (Table S1). Detailed lists of identified binding sites and probe clusters (Tables S2-4). Experimental results (MS, NMR) proving the labelling for the hits (Pages S6-S33).

## 1. RMSD calculations between STAT5B N-terminal domain structures

**Supplementary Table S1.**
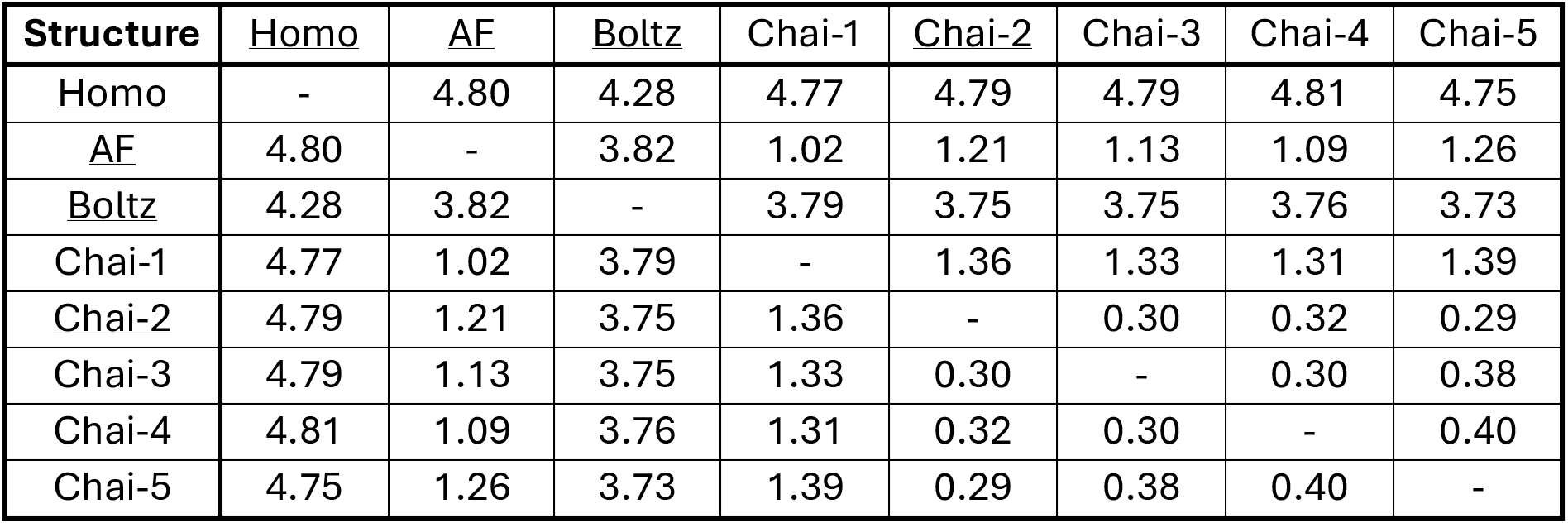
Pairwise backbone atom RMSD [Å] calculated between different STAT5B N-terminal domain structures (Homo = Homology, AF = AlphaFold model AF-P51692-F1). Used structures for this study are underlined.

| **Structure** | Homo | AF | Boltz | Chai-1 | <u>Chai-2</u> | Chai-3 | Chai-4 | Chai-5 |
| --- | --- | --- | --- | --- | --- | --- | --- | --- |
| <u>Homo</u> | - | 4.80 | 4.28 | 4.77 | 4.79 | 4.79 | 4.81 | 4.75 |
| <u>AF</u> | 4.80 | - | 3.82 | 1.02 | 1.21 | 1.13 | 1.09 | 1.26 |
| <u>Boltz</u> | 4.28 | 3.82 | - | 3.79 | 3.75 | 3.75 | 3.76 | 3.73 |
| Chai-1 | 4.77 | 1.02 | 3.79 | - | 1.36 | 1.33 | 1.31 | 1.39 |
| <u>Chai-2</u> | 4.79 | 1.21 | 3.75 | 1.36 | - | 0.30 | 0.32 | 0.29 |
| Chai-3 | 4.79 | 1.13 | 3.75 | 1.33 | 0.30 | - | 0.30 | 0.38 |
| Chai-4 | 4.81 | 1.09 | 3.76 | 1.31 | 0.32 | 0.30 | - | 0.40 |
| Chai-5 | 4.75 | 1.26 | 3.73 | 1.39 | 0.29 | 0.38 | 0.40 | - |

**Supplementary Table S2.** Number of probe molecules found in each cluster on KRAS, and which known binding site, identified site from computational studies or experimentally identified site/hit the appropriate cluster have found.

| Open clusters |  |  |  |  |
| --- | --- | --- | --- | --- |
| Cluster ID | Number of probe molecules | Corresponding known binding site(s)<sup>1-3</sup> | Corresponding identified site(s) from computational studies<sup>4-6</sup> | Corresponding experimentally identified hit(s)<sup>1,7-10</sup> |
| O1 | 2 | SWI/SWII pocket | Xe2<sup>5</sup><br>p1 site<sup>6</sup> | - |
| O2 | 2 | GDP binding site | - | Cluster 4<sup>10</sup> |
| O3 | 4 | Dimer interface | - | PhP060<sup>1</sup><br>F5, F8 |
| O4 | 5 | - | - | DARPin<sup>7</sup> |
| O5 | 9 | Calmodulin binding interface<sup>2,3</sup> | P3 site<sup>4</sup><br>p3 site<sup>6</sup> | - |
| O6 | 10 | Dimer interface | P7 site<sup>4</sup><br>Xe7<sup>5</sup> | PhP060<sup>1</sup><br>F5, F8 |
| O7 | 14 | Switch-II pocket | P2 site<sup>4</sup><br>Xe1<sup>5</sup><br>p2 site<sup>6</sup> | Adagrasib<sup>8</sup><br>Sotorasib<sup>9</sup><br>PhP072<sup>1</sup><br>F1, F3, F6<br>Cluster 1<sup>10</sup> |
| O8 | 18 | SWI/SWII pocket<br>~ Switch-II loop | P1 site<sup>4</sup><br>Xe2<sup>5</sup><br>p1 site<sup>6</sup> | PhP072<sup>1</sup><br>F7 |
| O9 | 31 | GDP binding site<br>~Switch-I loop | - | Cluster 6, Cluster 8<sup>10</sup> |
| Cryptic clusters |  |  |  |  |
| Cluster ID | Number of probe molecules | Corresponding known binding site(s)<sup>1-3</sup> | Corresponding identified site(s) from computational studies<sup>4-6</sup> | Corresponding experimentally identified hit(s)<sup>1,7-10</sup> |
| C1 | 2 | Switch-II loop | Xe4<sup>5</sup> | - |
| C2 | 3 | - | Xe3<sup>5</sup><br>p3 site<sup>6</sup> | DARPin<sup>7</sup><br>F1, F3, F4, F9 |
| C3 | 4 | RAS-binding domain | P4 site<sup>4</sup> | PhP060, PhP072<sup>1</sup><br>F2<br>Cluster 3<sup>10</sup> |
| C4 | 11 | Dimer interface | Xe6<sup>5</sup> | PhP071<sup>1</sup><br>F8 |
| C5 | 12 | - | - | - |
| C6 | 12 | - | - | F2, F4, F5, F10 |
| C7 | 19 | - | - | F2, F7, F10 |
| C8 | 20 | - | P8 site<sup>4</sup><br>p3 site<sup>6</sup> | DARPin<sup>7</sup><br>F1, F3, F4, F9<br>Cluster 2<sup>10</sup> |
| C9 | 26 | SWI/SWII pocket | - | F2, F7 |
| C10 | 44 | RAS-binding domain | P4 site<sup>4</sup><br>p4 site<sup>6</sup> | PhP060, PhP072<sup>1</sup> |
| C11 | 96 | - | P6 site<sup>4</sup> | F2, F4, F5, F6 |
| C12 | 474 | - | P5 site<sup>4</sup><br>Xe8<sup>5</sup><br>p5 site<sup>6</sup> | Cluster 7<sup>10</sup> |

**Supplementary Table S3.** Number of probe molecules found in each cluster on each STAT5B NTD structure.

| Open clusters |  |  |  |  |  |  |  |
| --- | --- | --- | --- | --- | --- | --- | --- |
| Homology model | Number of probe molecules | AlphaFold model | Number of probe molecules | Boltz model | Number of probe molecules | Chai model | Number of probe molecules |
| H-O1 | 2 | A-O1 | 1 | B-O1 | 3 | C-O1 | 2 |
| H-O2 | 2 | A-O2 | 2 | B-O2 | 5 | C-O2 | 2 |
| H-O3 | 2 | A-O3 | 3 | B-O3 | 5 | C-O3 | 7 |
| H-O4 | 2 | A-O4 | 5 | B-O4 | 12 | C-O4 | 8 |
| H-O5 | 5 | A-O5 | 6 | B-O5 | 20 | C-O5 | 22 |
| H-O6 | 17 | A-O6 | 12 | B-O6 | 21 | C-O6 | 26 |
| H-O7 | 19 | A-O7 | 16 | B-O7 | 27 | C-O7 | 28 |
| H-O8 | 44 | A-O8 | 22 |  |  |  |  |
|  |  | A-O9 | 26 |  |  |  |  |
| Cryptic clusters |  |  |  |  |  |  |  |
| Homology model | Number of probe molecules | AlphaFold model | Number of probe molecules | Boltz model | Number of probe molecules | Chai model | Number of probe molecules |
| H-C1 | 2 | A-C1 | 2 | B-C1 | 2 | C-C1 | 1 |
| H-C2 | 4 | A-C2 | 17 | B-C2 | 4 | C-C2 | 1 |
| H-C3 | 1157 | A-C3 | 116 | B-C3 | 6 | C-C3 | 17 |
| H-C4 | 1196 | A-C4 | 190 | B-C4 | 48 | C-C4 | 26 |
|  |  | A-C5 | 449 | B-C5 | 81 | C-C5 | 59 |
|  |  | A-C6 | 828 |  |  | C-C6 | 69 |
|  |  |  |  |  |  | C-C7 | 612 |
|  |  |  |  |  |  | C-C8 | 779 |

**Supplementary Table S4.** Locationally corresponding other clusters, known binding interfaces or experimentally identified hits to identified clusters. Cluster used as location reference is framed. Only those clusters were treated as location reference, whose were not correlating to other clusters in previous rows.

| Open cluster used as location reference |  |  |  |  |  |
| --- | --- | --- | --- | --- | --- |
| Homology model | AlphaFold model | Boltz model | Chai model | Corresponding known binding interface | Corresponding experimentally identified hit(s)<sup>1</sup> |
| **H-O1** | A-O6 | B-O2 | C-O4 | - | - |
| **H-O2** | - | B-C5 | C-O3 | - | - |
| **H-O3** | A-C5 | - | C-C5 | - | - |
| **H-O4** | A-O1<br>A-O4 | B-O3 | C-O2 | - | PhP065<sup>1</sup> |
| **H-O5** | A-O8 | B-O5 | C-O6 | Handshake interface | L1 |
| **H-O6** | A-O8 | B-O5 | C-O6 | Handshake interface | - |
| **H-O7** | A-C6 | B-O1 | C-C7 | Ni<sup>2+</sup> – mediated interface | L1 |
| **H-O8** | A-O9 | B-O7 | C-O7 | - | L1 |
| - | **A-O2** | - | - | Ni<sup>2+</sup> – mediated interface | L1<br>L2 |
| - | **A-O3** | - | C-O3 | - | - |
| - | **A-O5** | B-O6 | C-O5 | - | - |
| - | **A-O7** | B-O6 | C-O5 | - | - |
| - | A-C3 | **B-O4** | C-C4 | - | - |
| - | - | - | **C-O1** | - | - |
| Covib cluster used as location reference |  |  |  |  |  |
| Homology model cluster | AlphaFold model cluster | Boltz model cluster | Chai model cluster | Corresponding known binding interface | Corresponding experimentally identified hit(s) |
| **H-C1** | - | - | - | - | - |
| **H-C2** | - | - | C-C1<br>C-C2 | - | - |
| **H-C3** | - | - | C-C6 | - | PhP097<sup>1</sup><br>L1<br>L2 |
| **H-C4** | A-C4 | - | C-C8 | - | - |
| - | **A-C1** | - | - | - | - |
| - | **A-C2** | - | - | - | - |
| - | A-C6 | **B-C1** | - | Ni<sup>2+</sup> – mediated interface | L2 |
| - | - | **B-C2** | C-C3 | - | - |
| - | - | **B-C3** | - | - | - |
| - | - | **B-C4** | - | - | - |

## 2. Identified binding sites and probe clusters

## 3. Fragment hits labelling KRAS^G12D^

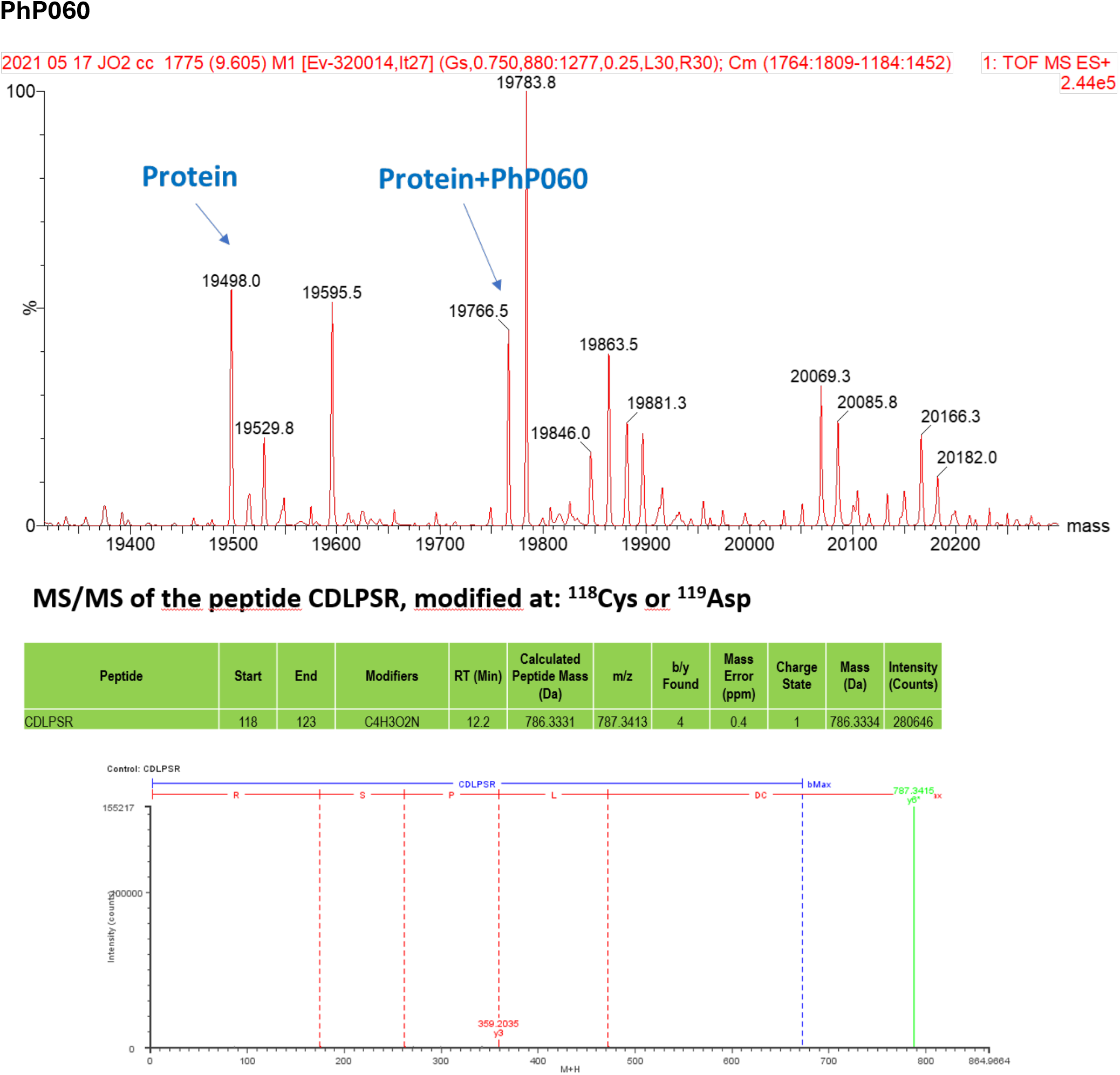
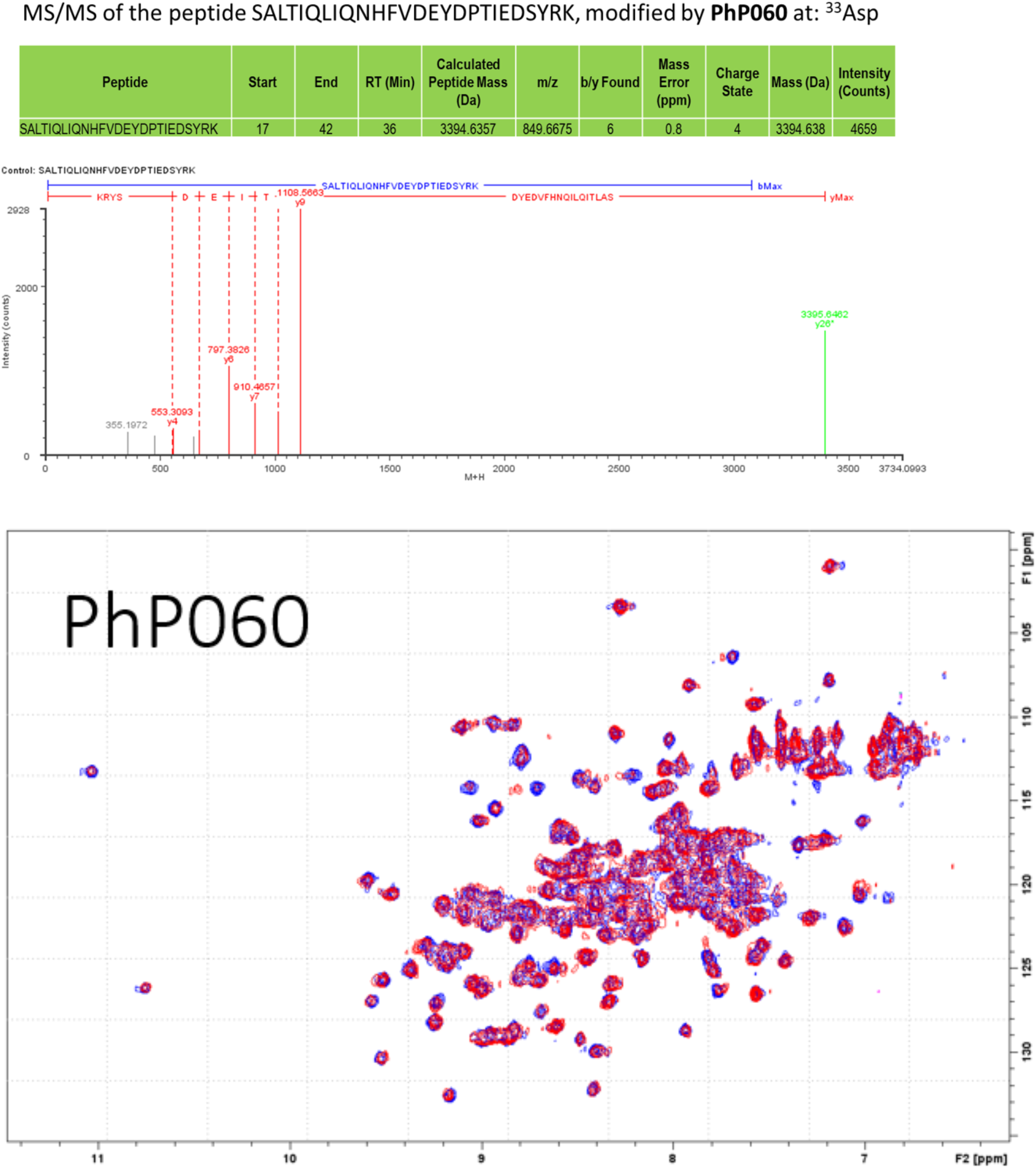
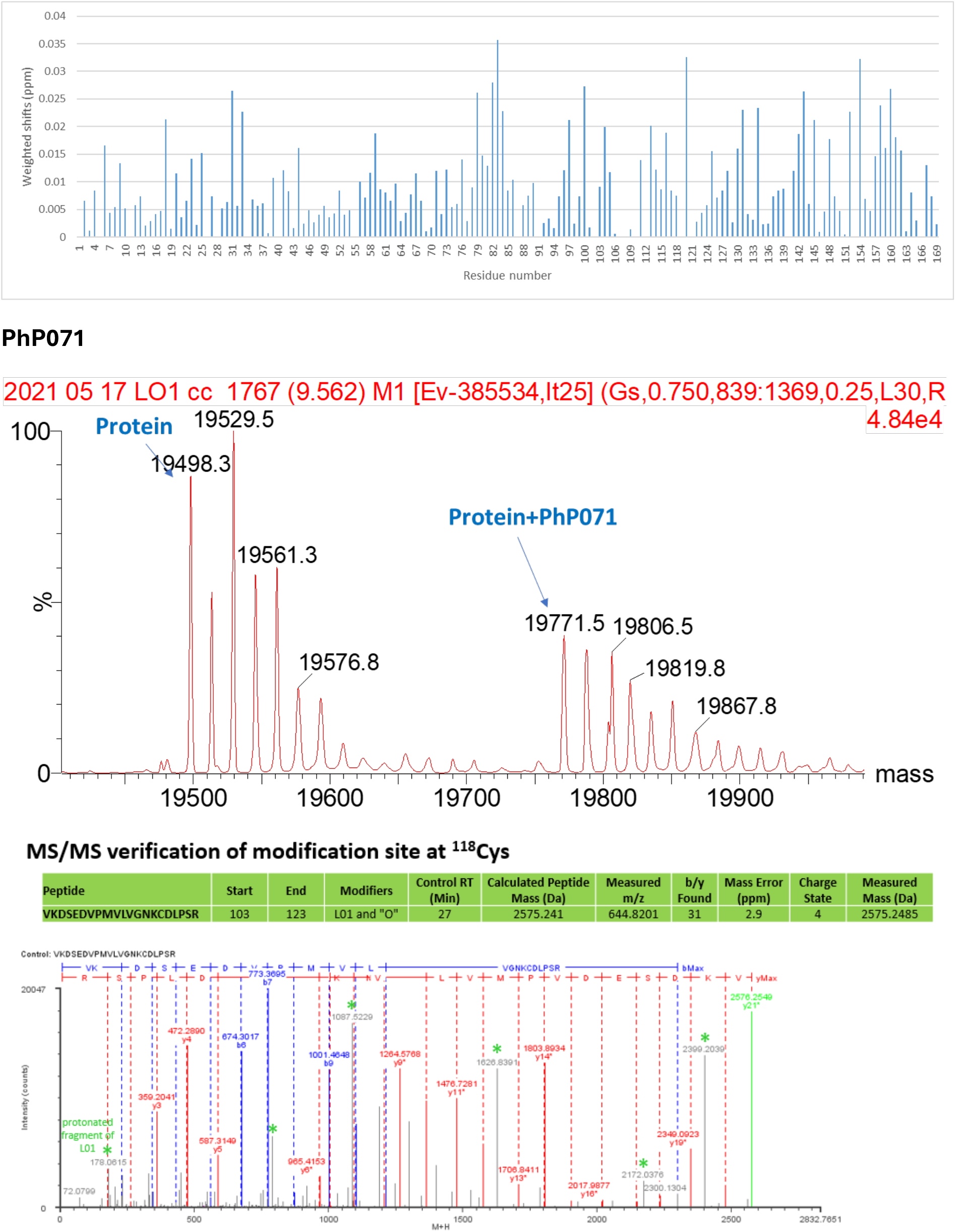
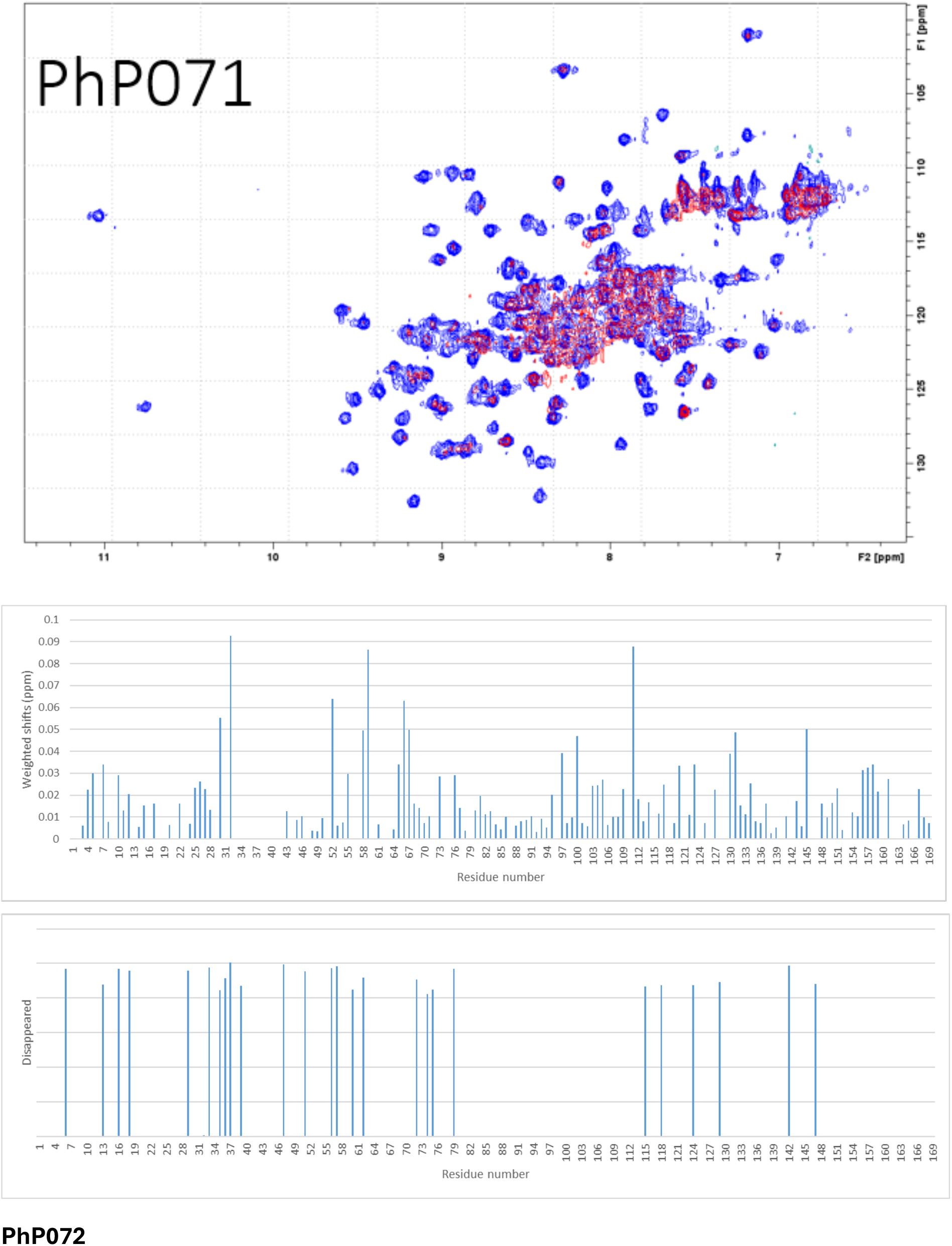
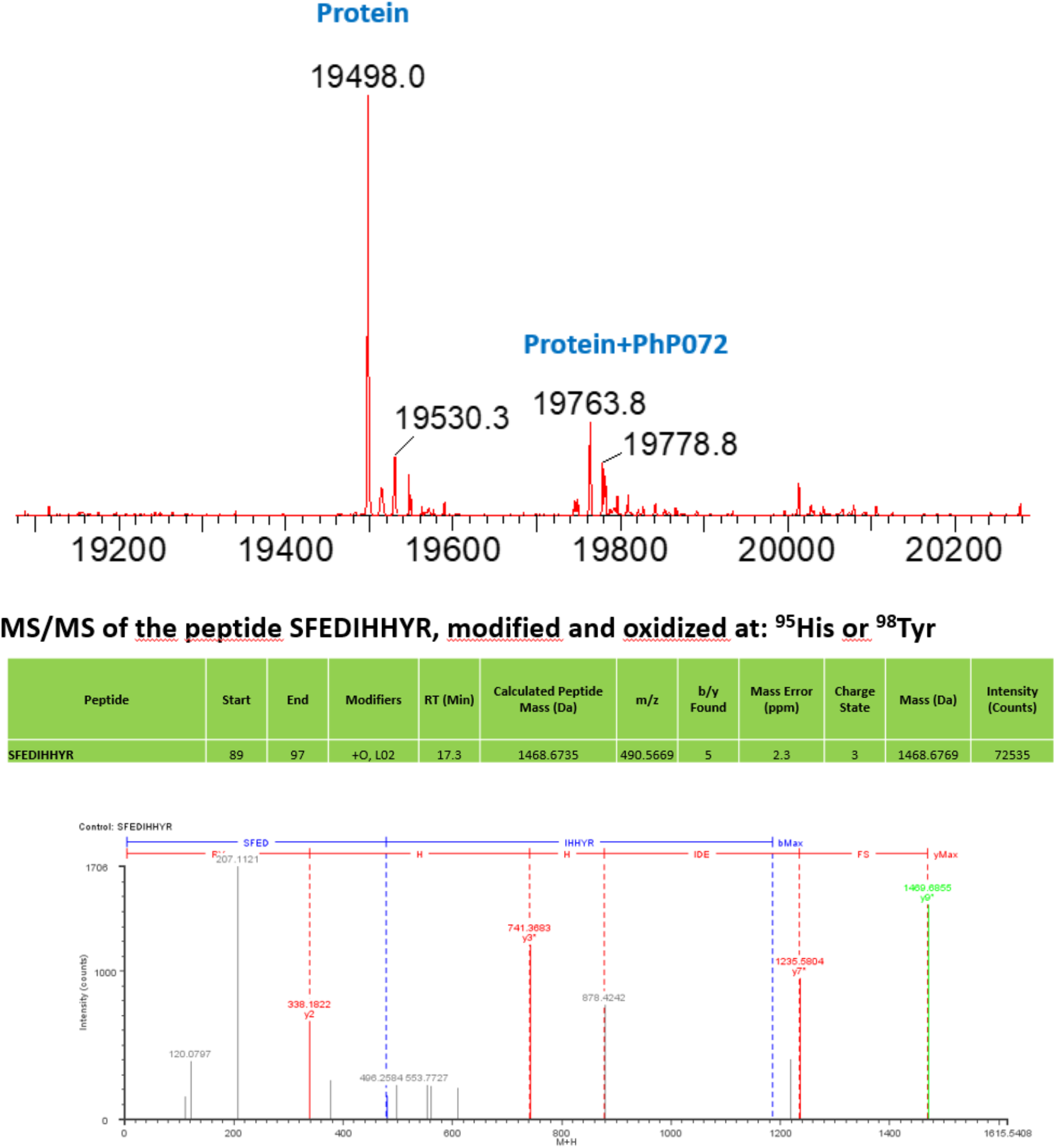
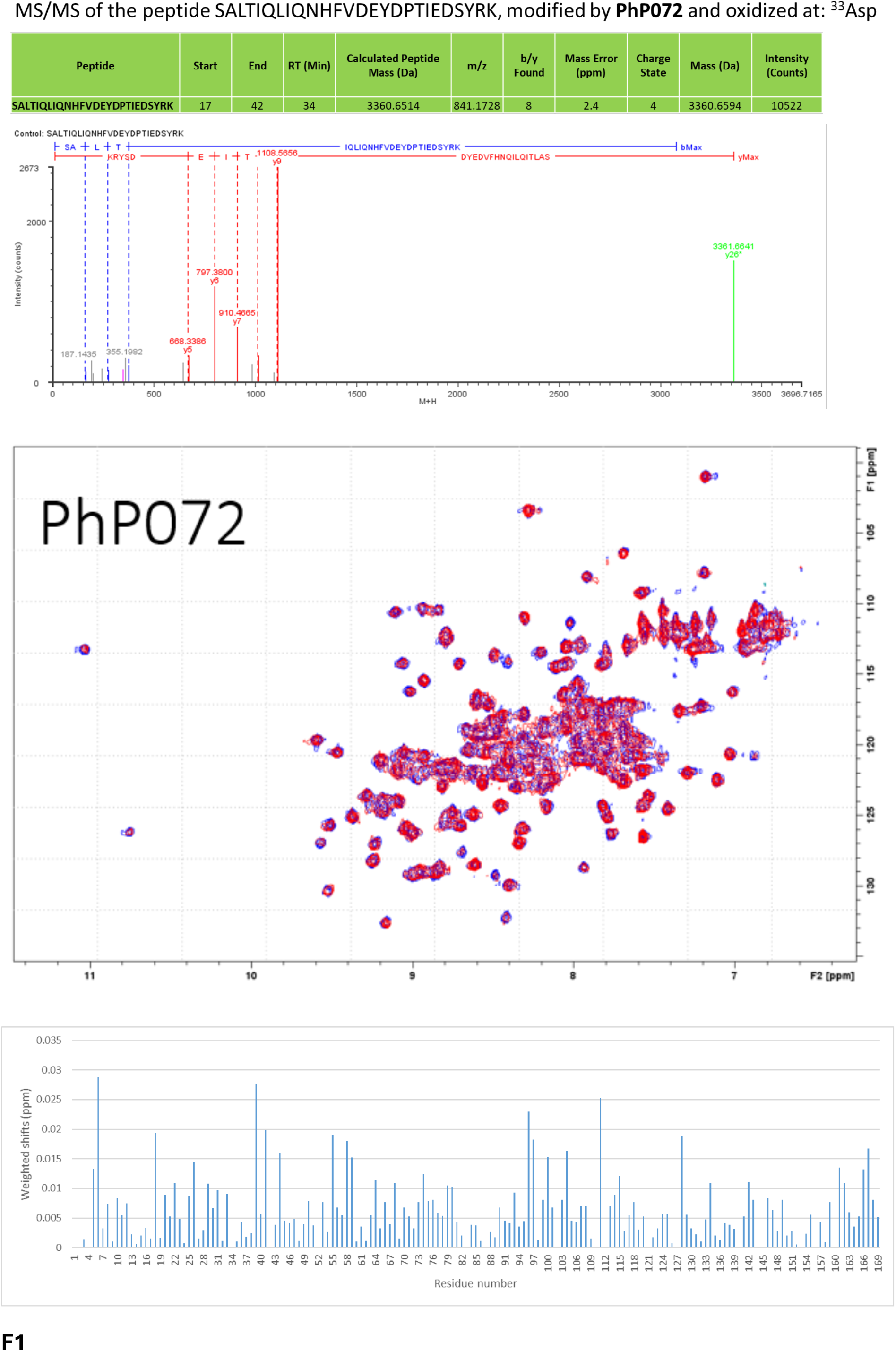
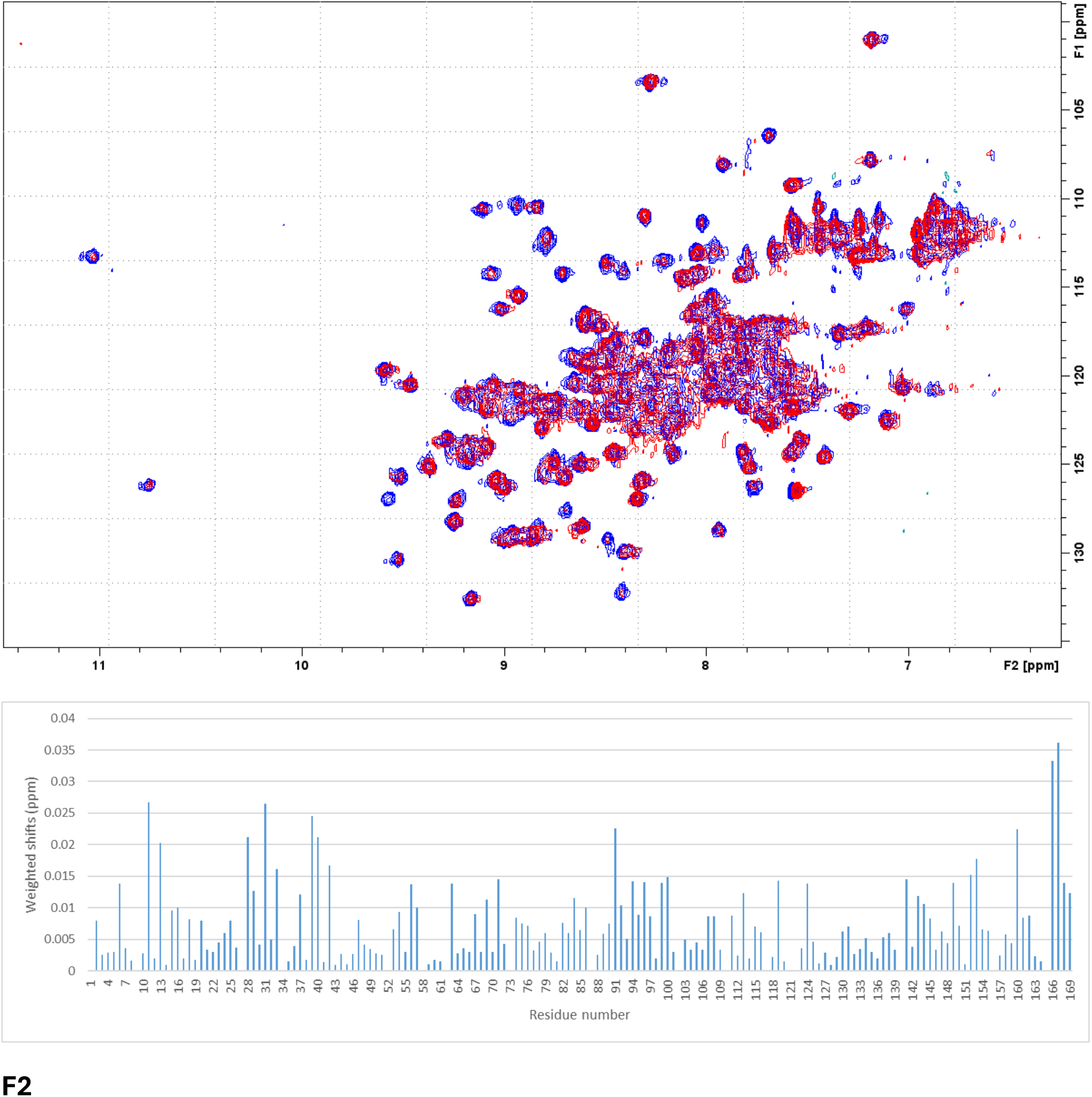
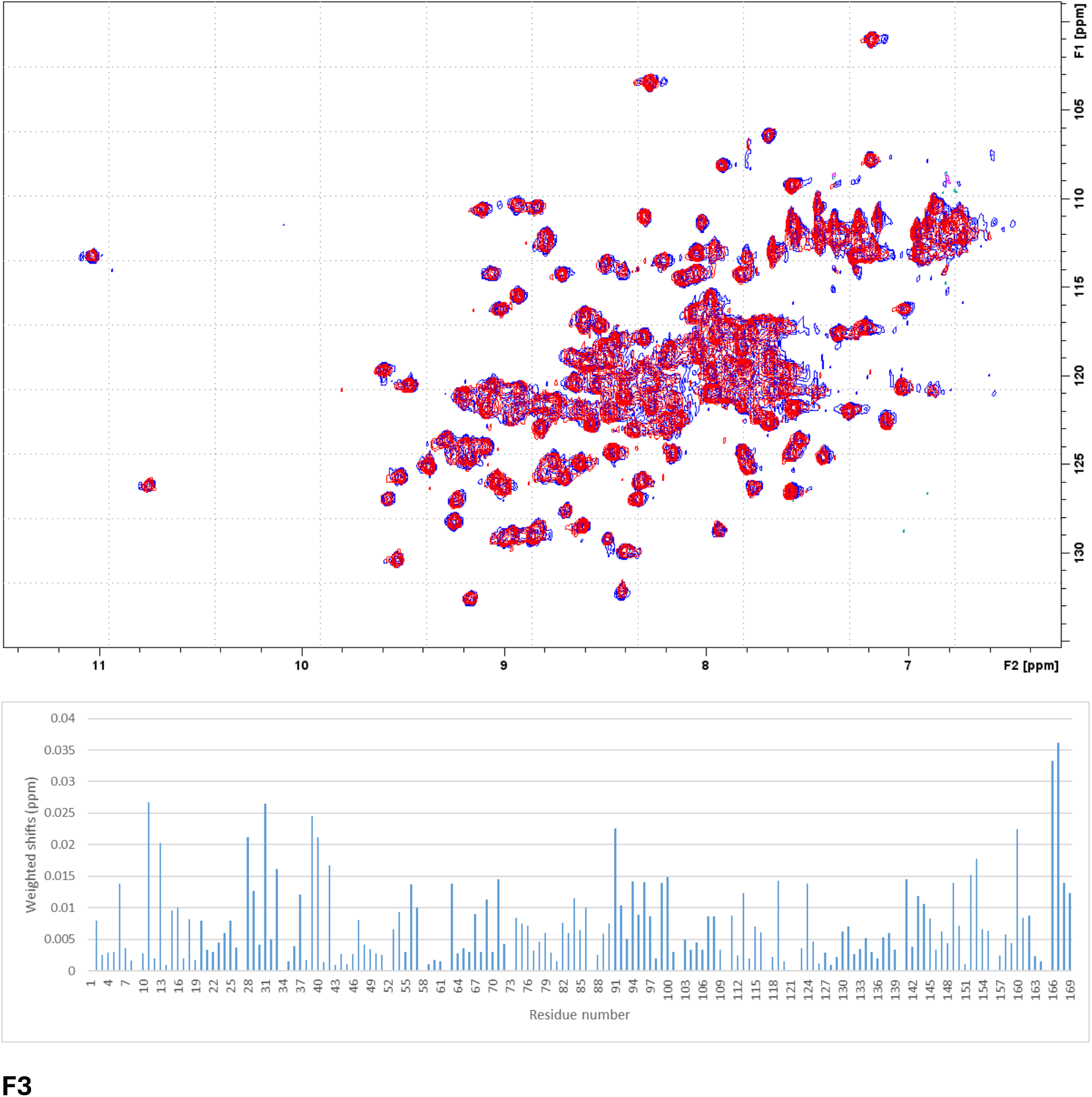
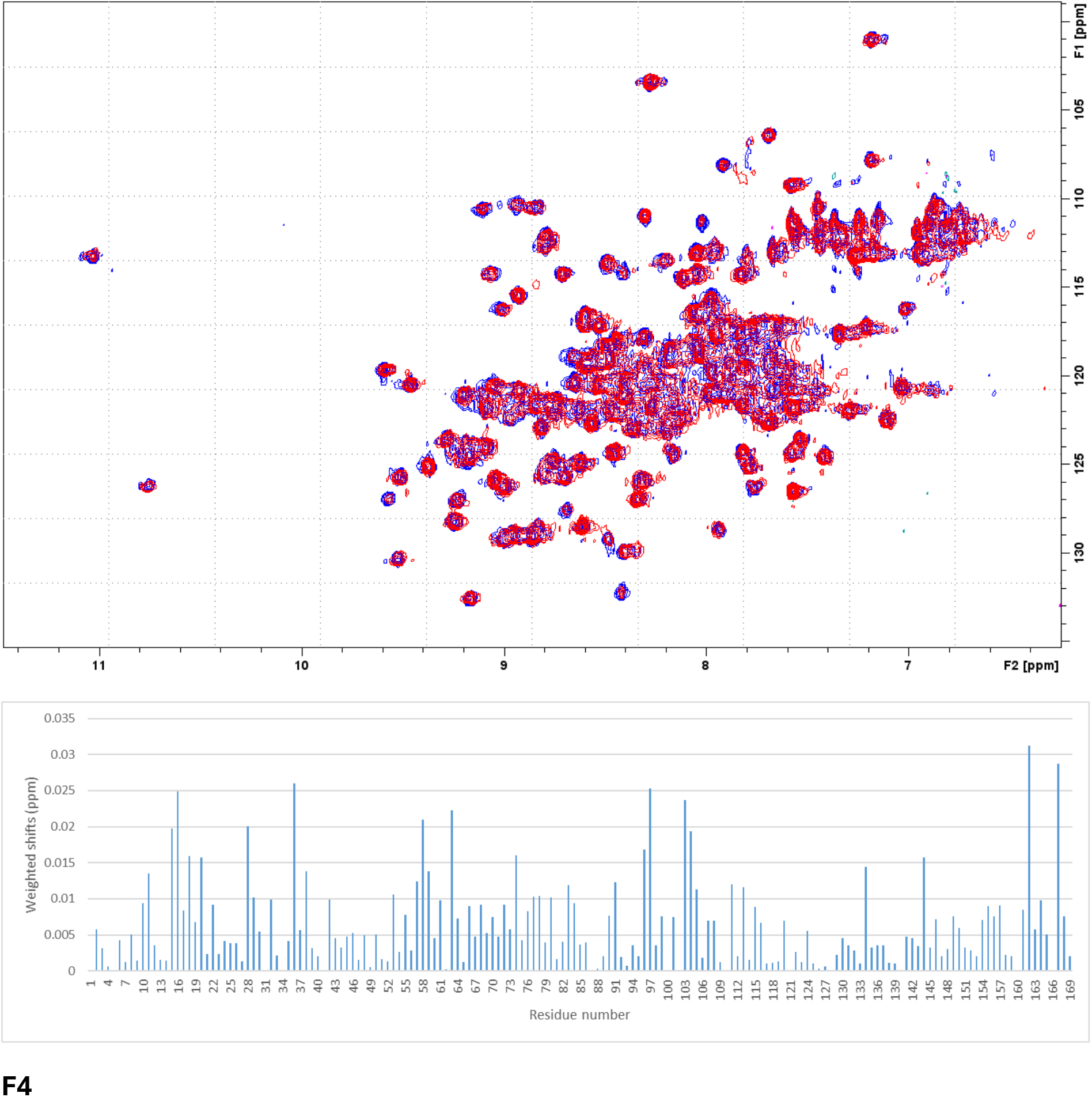
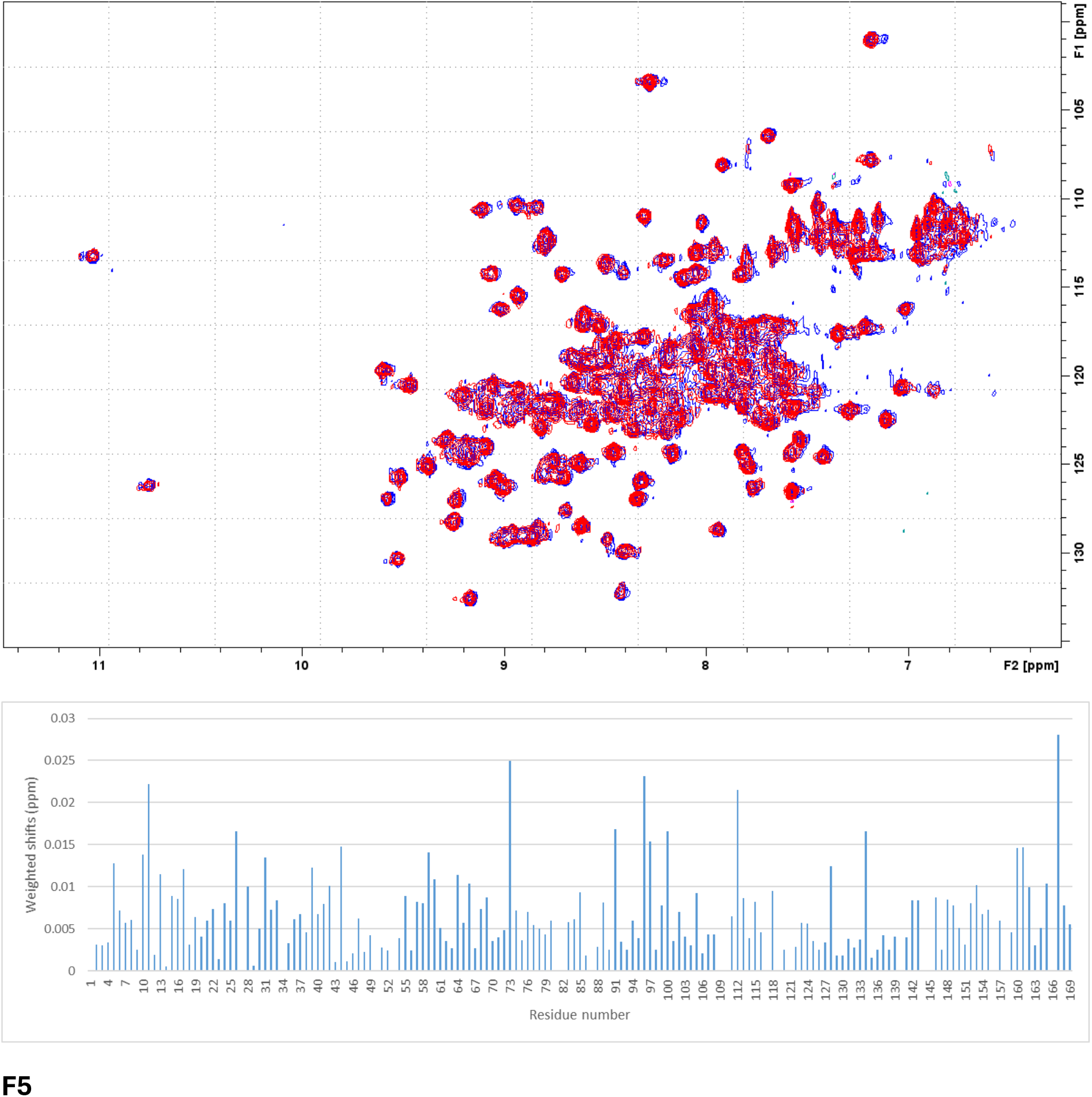
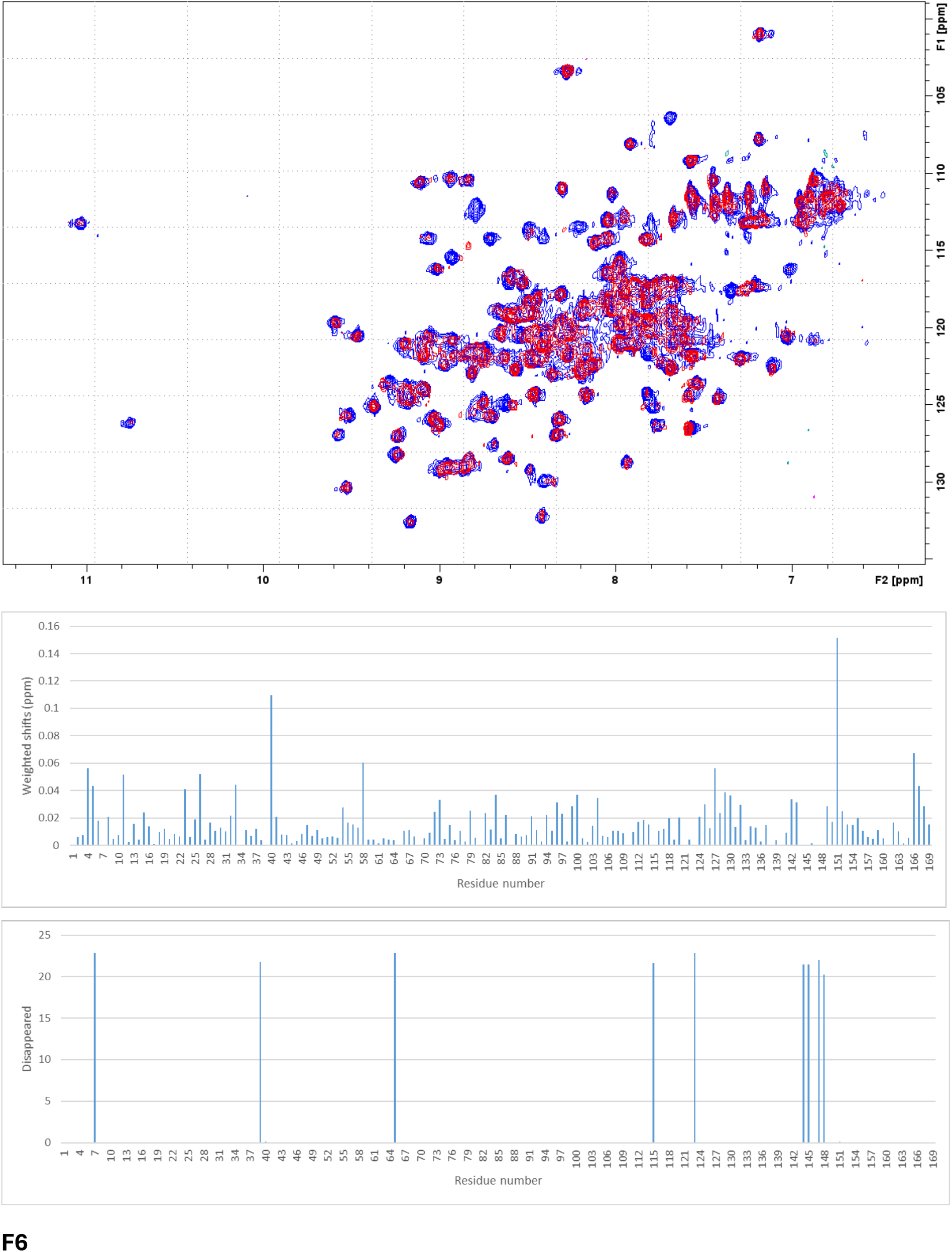
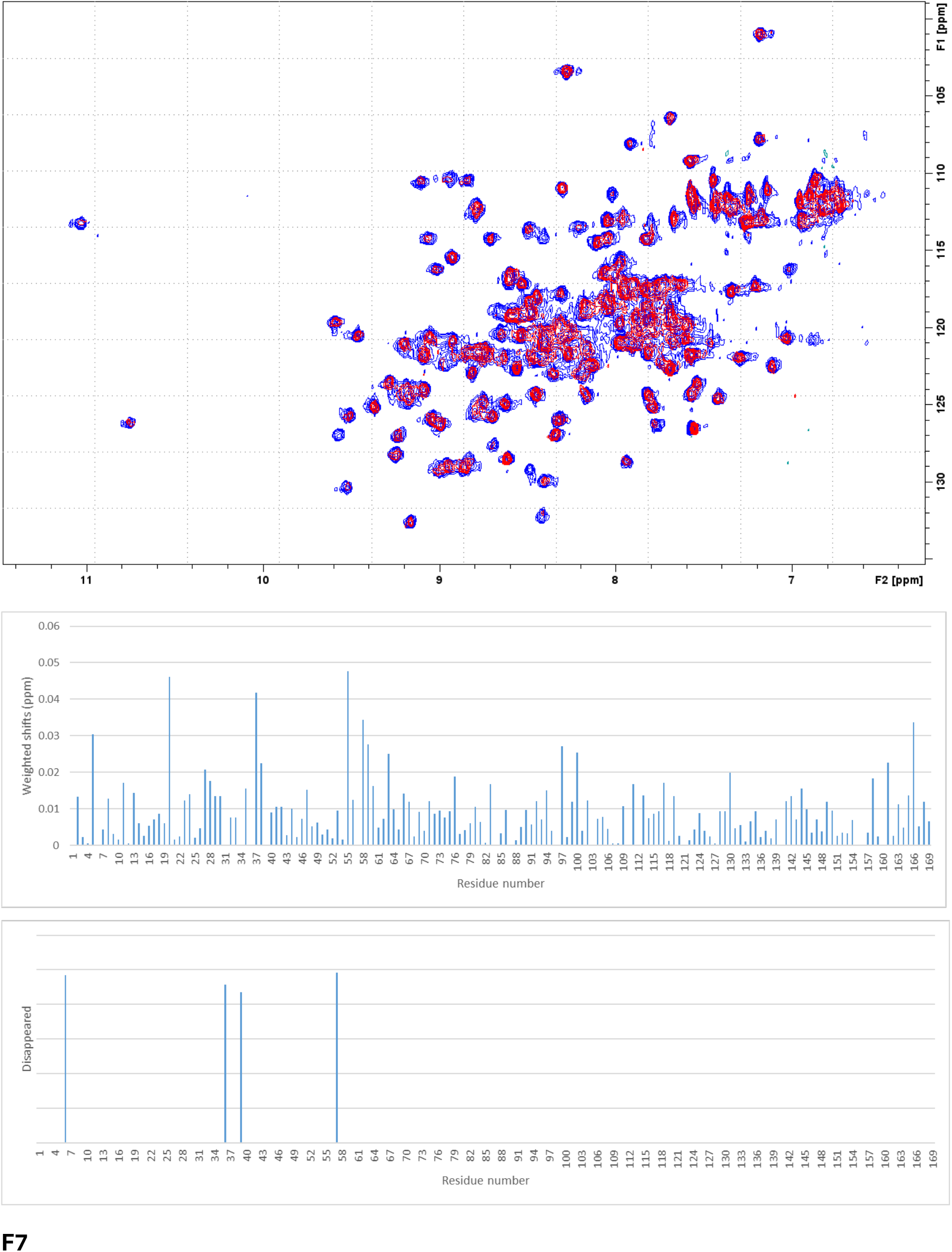
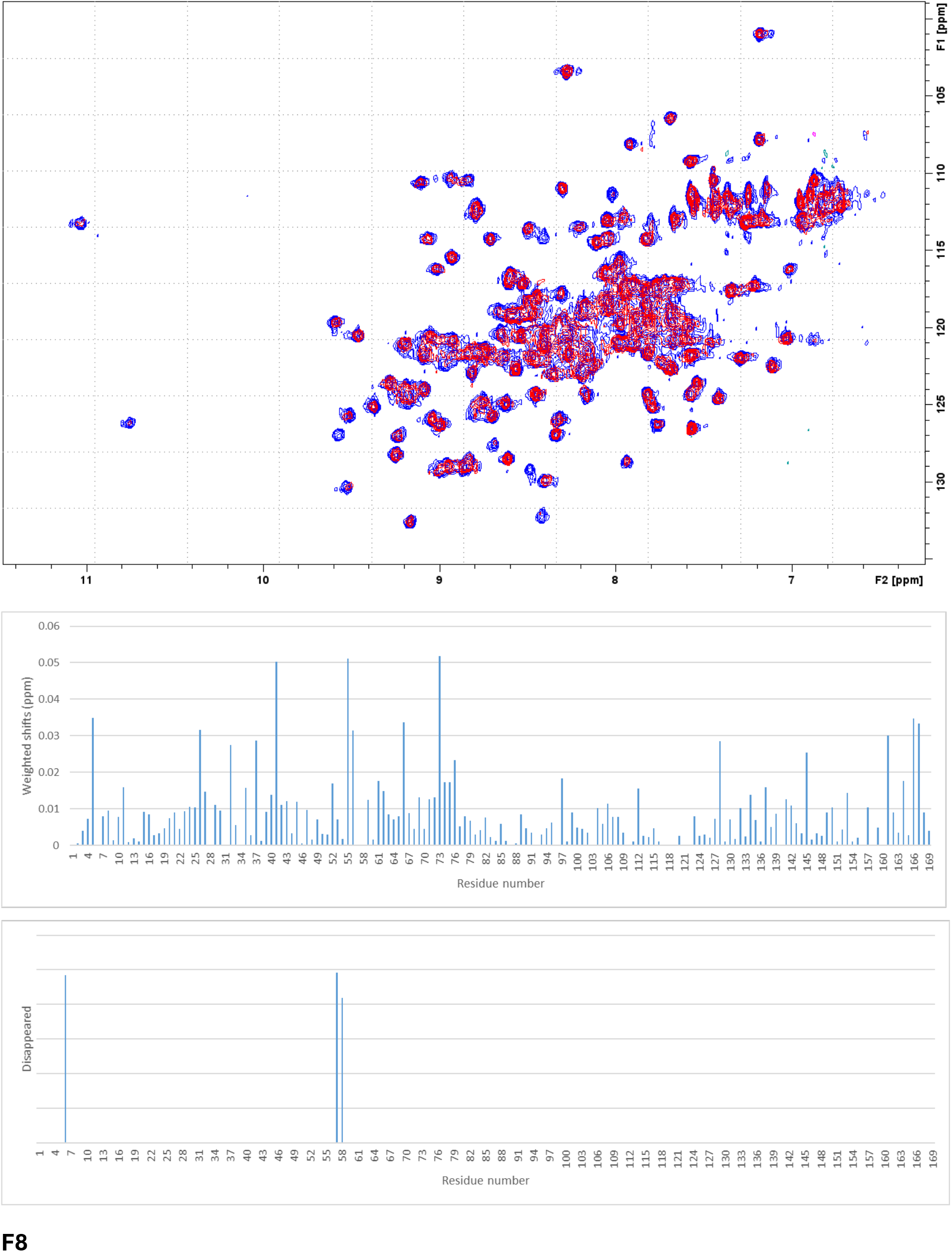
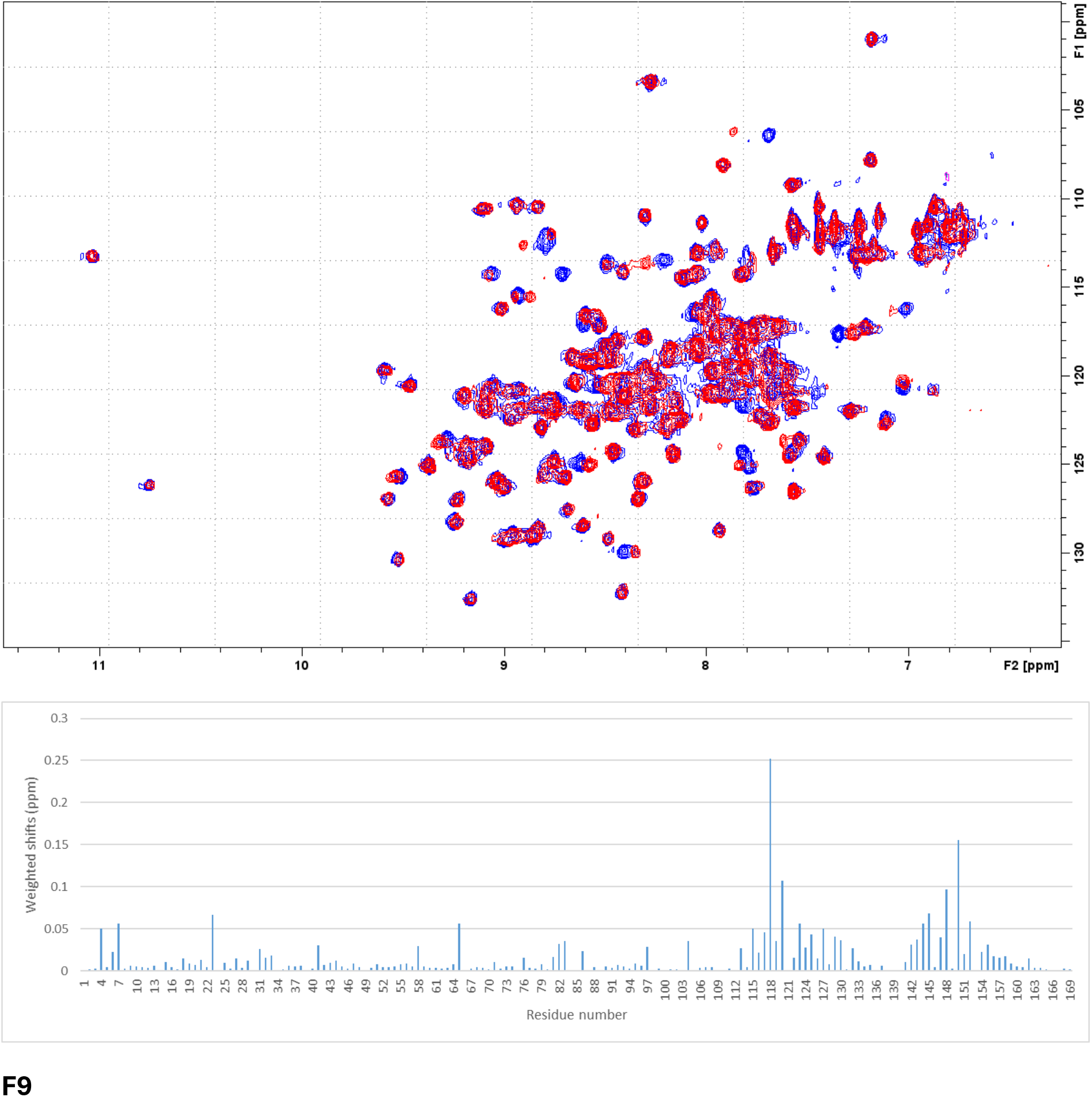
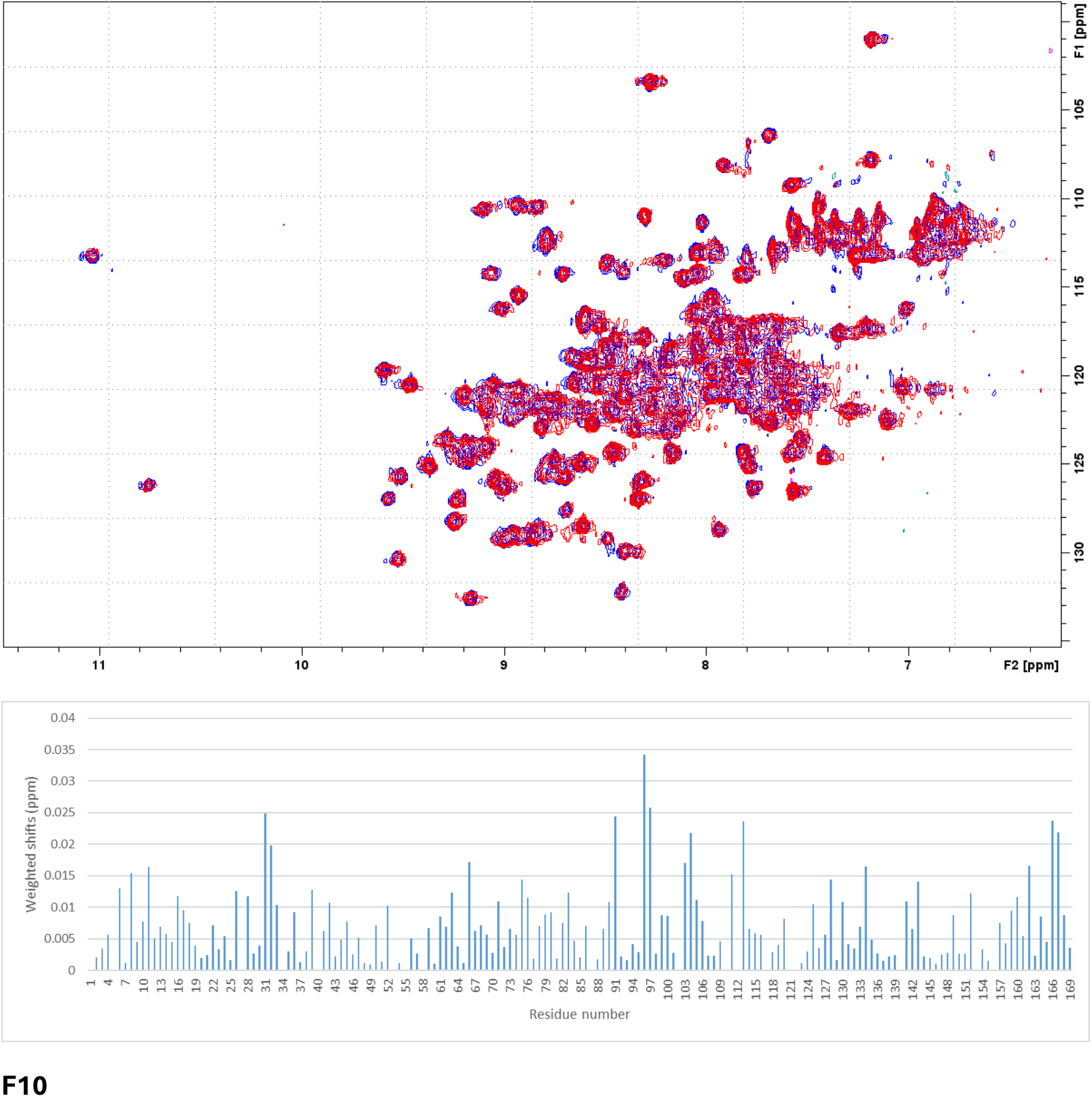
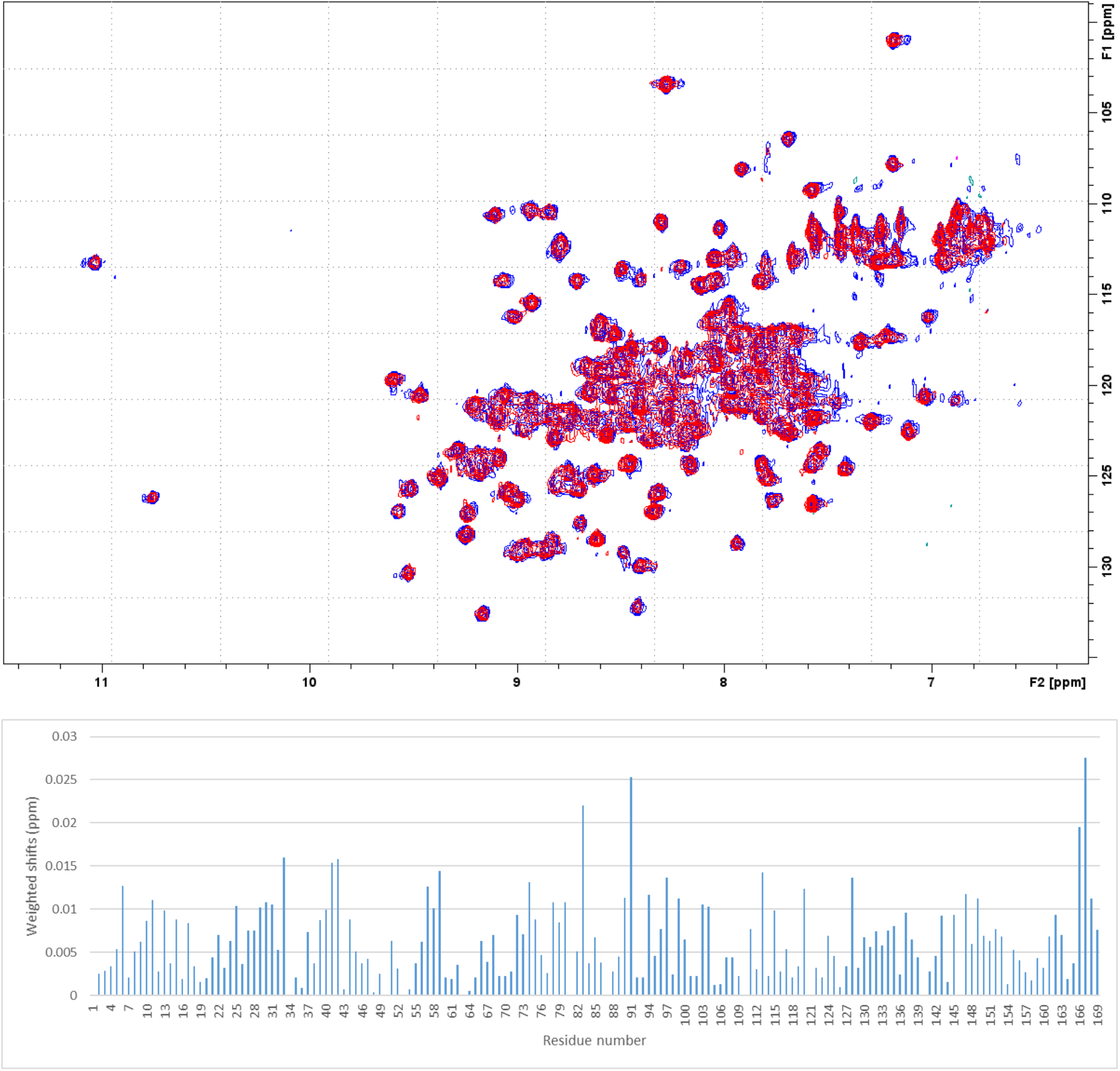

## 4. Fragment hits labelling STAT5B-NTD

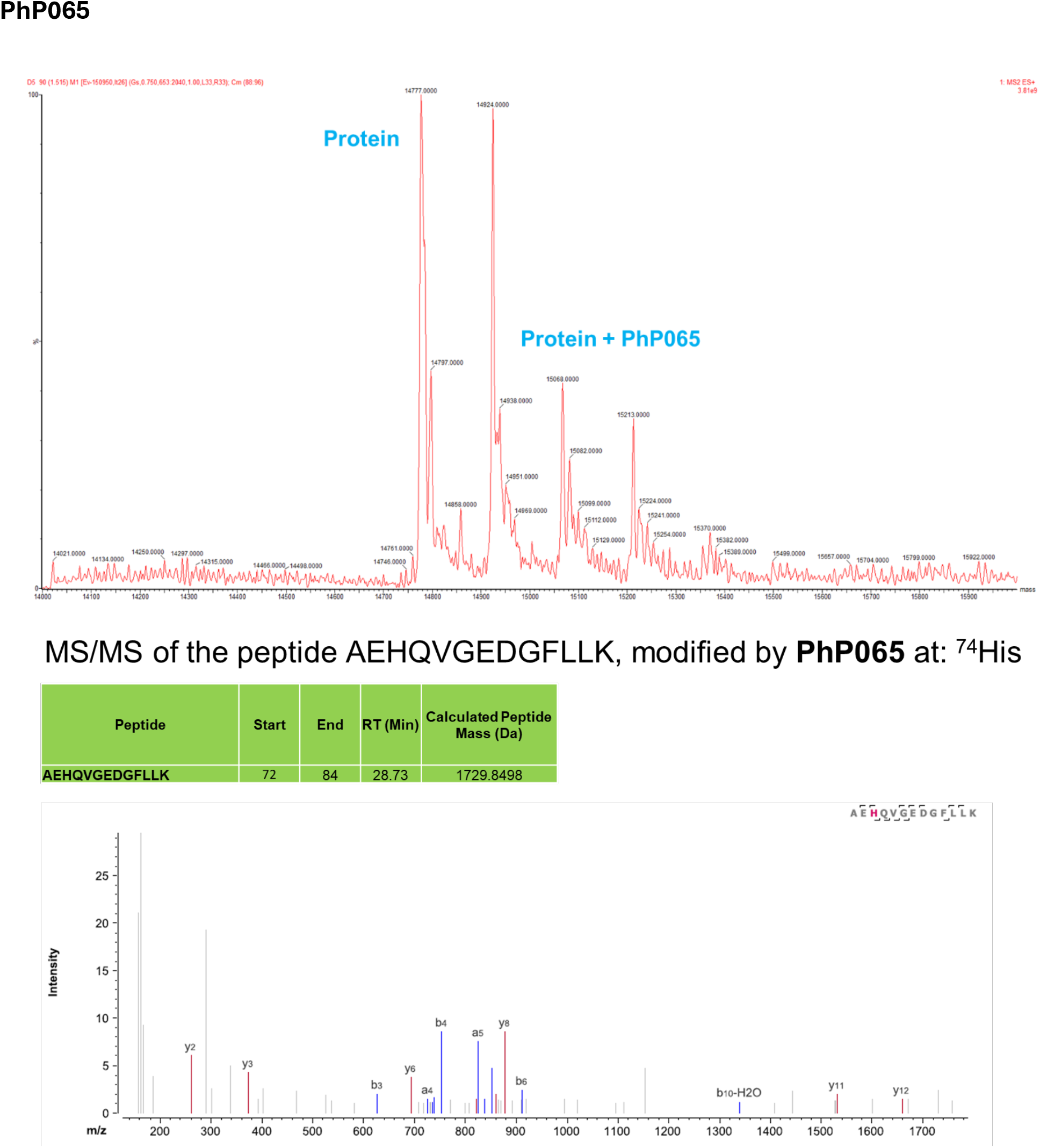
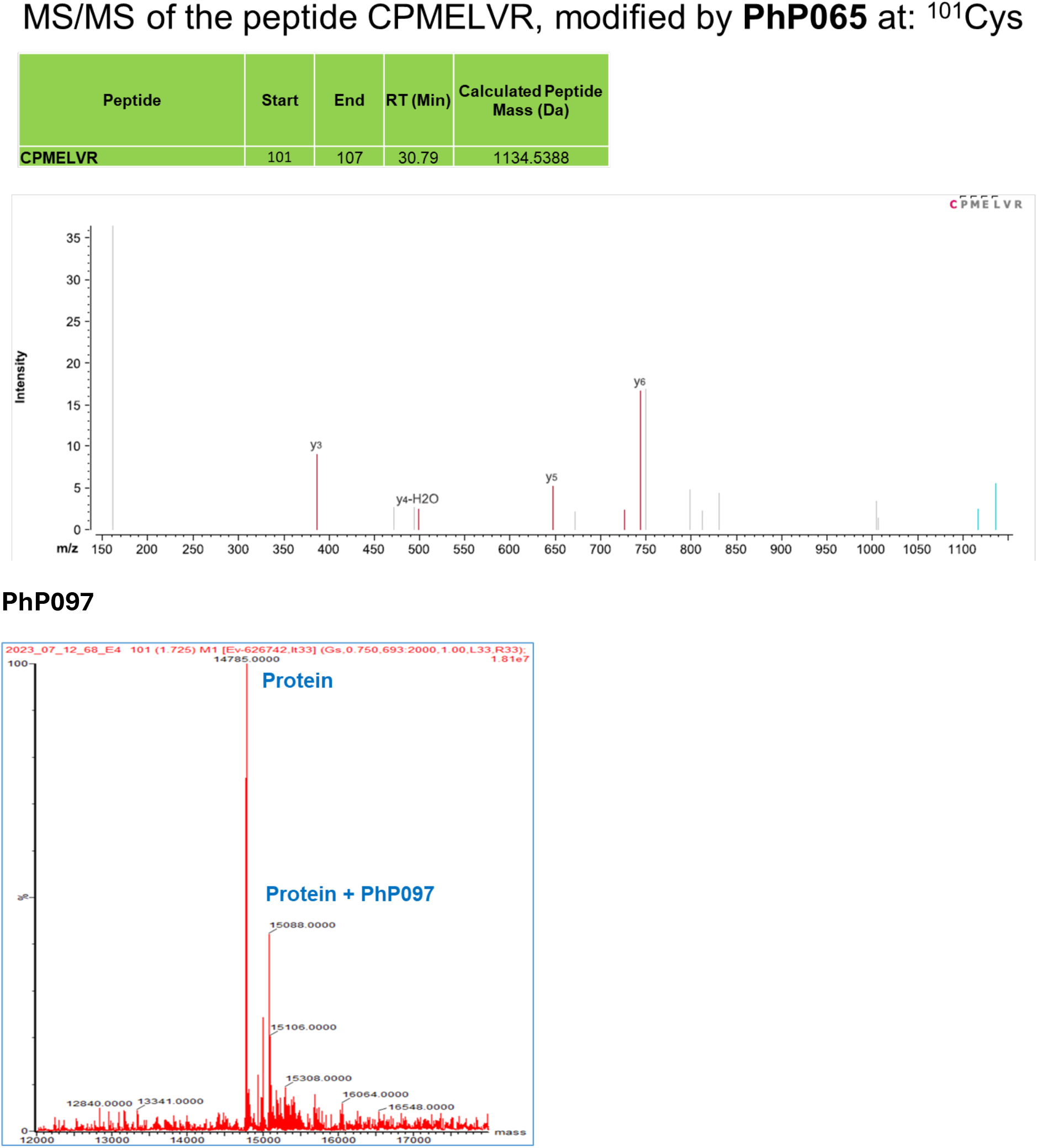
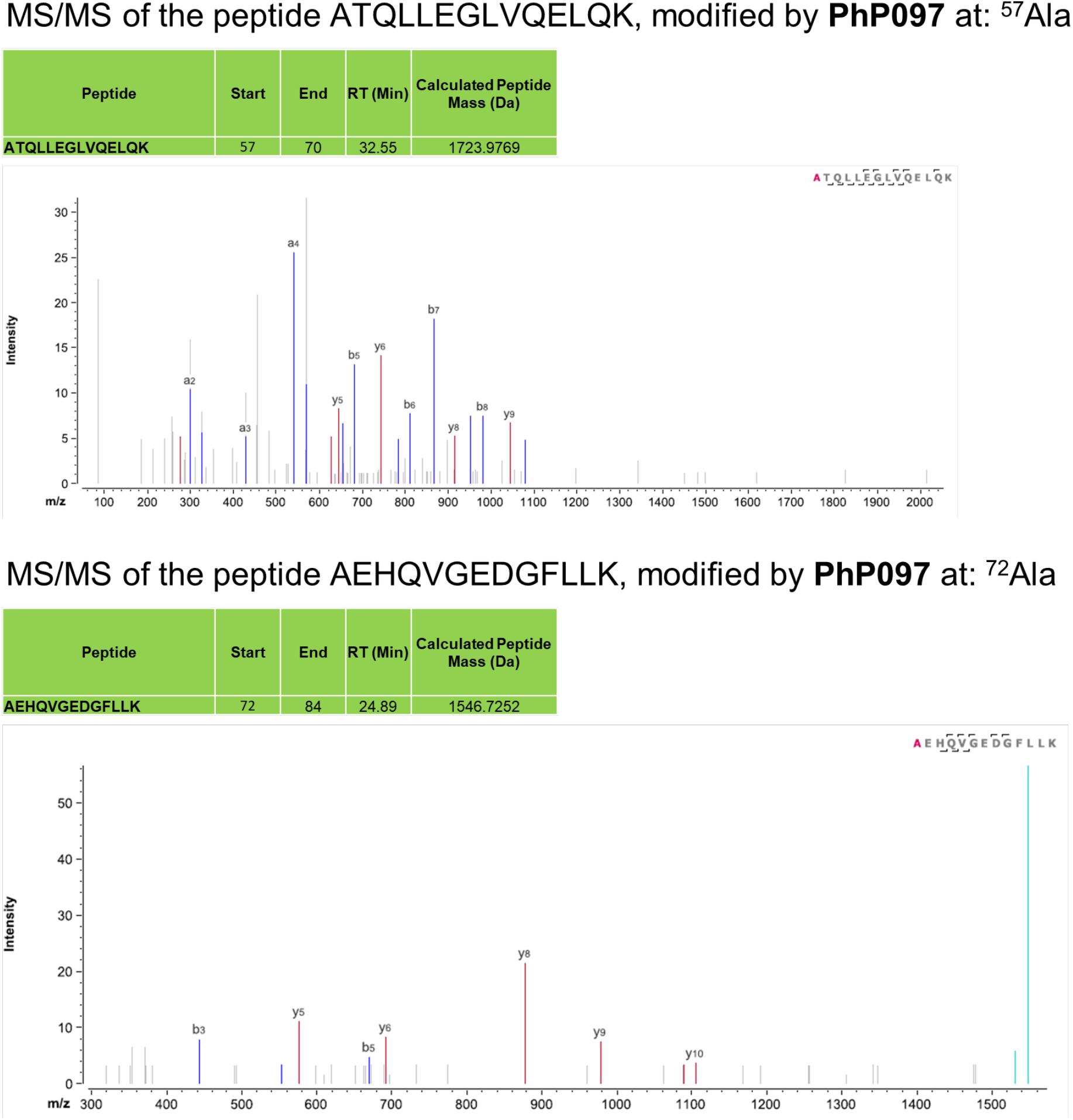
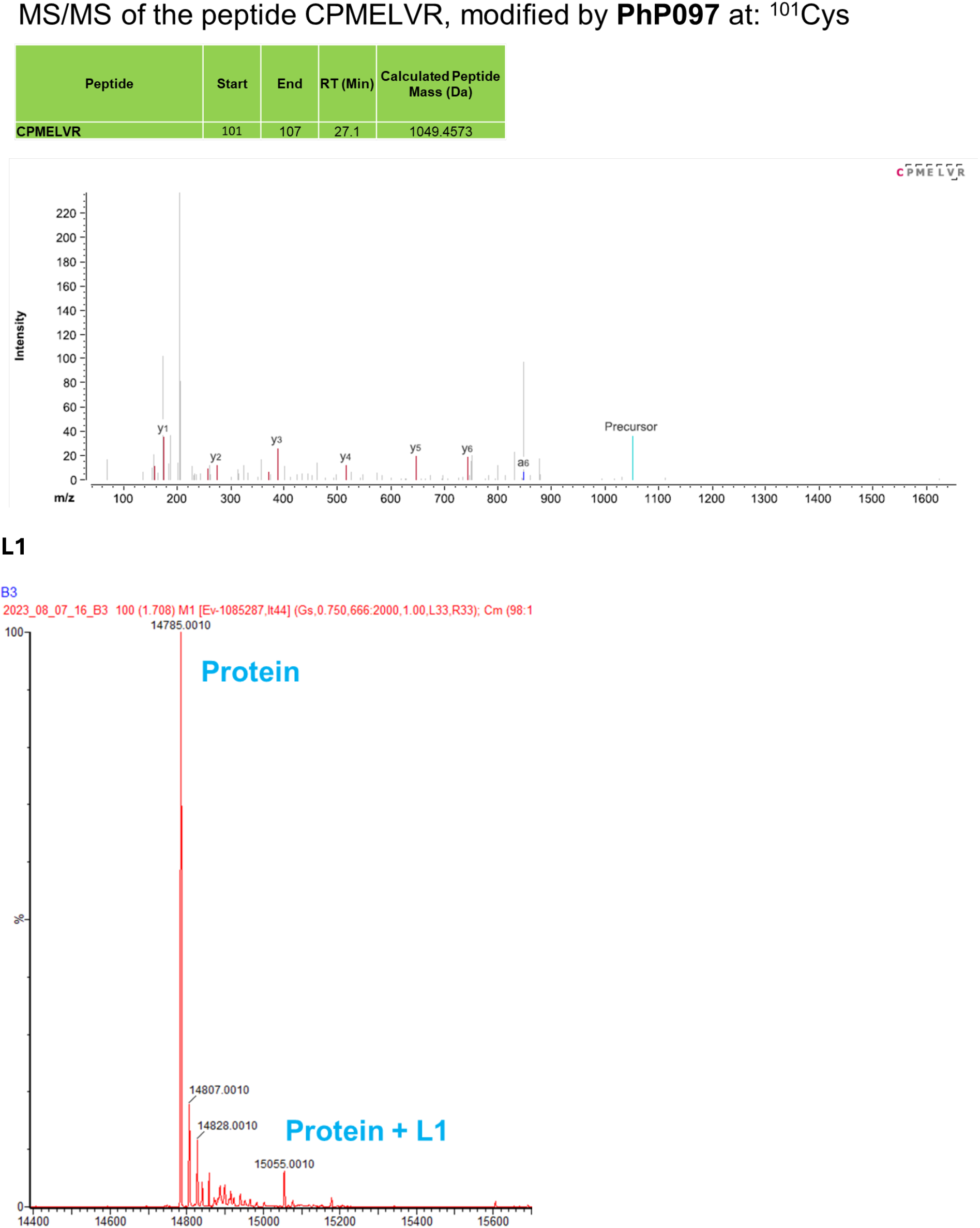
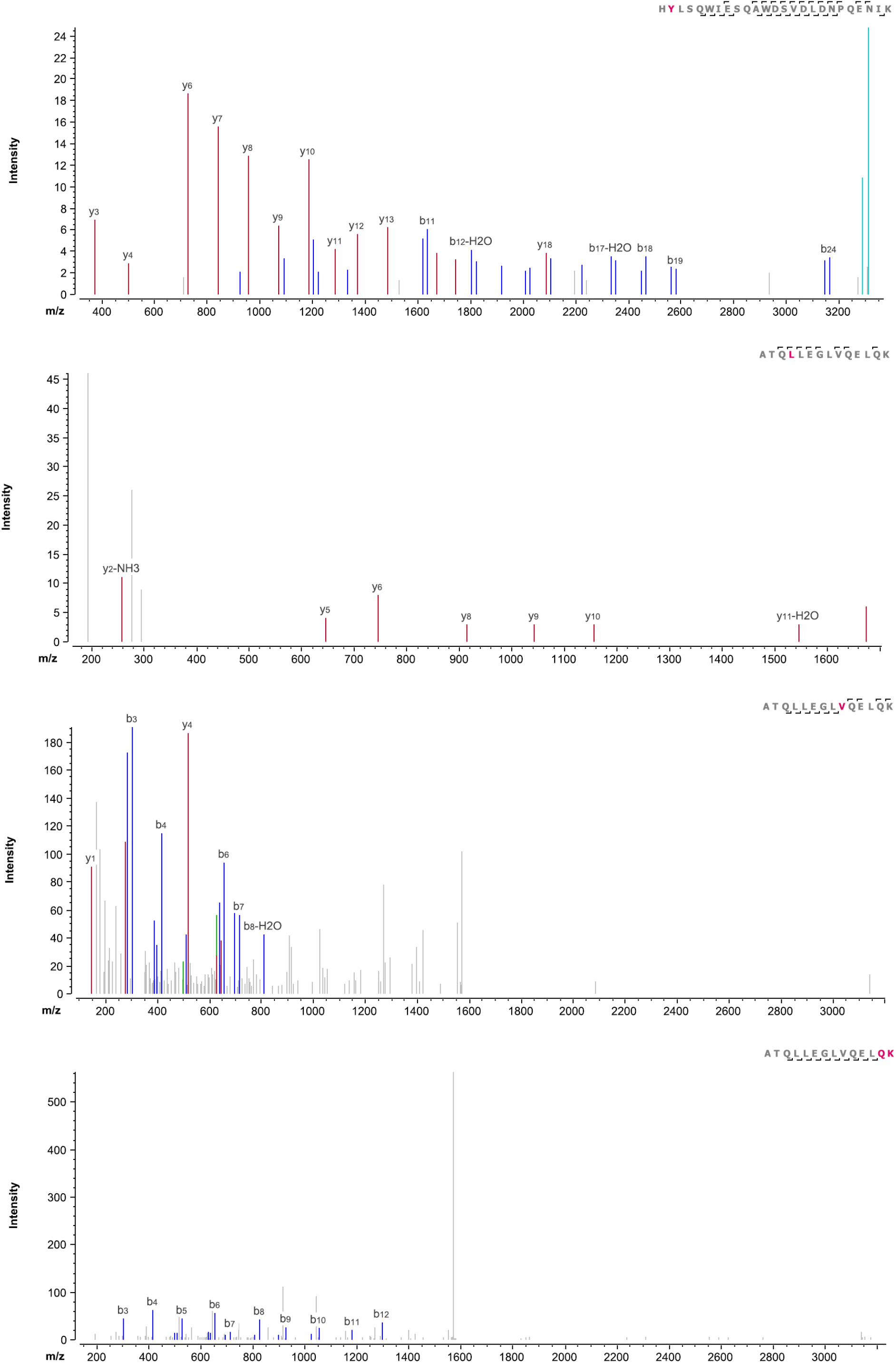
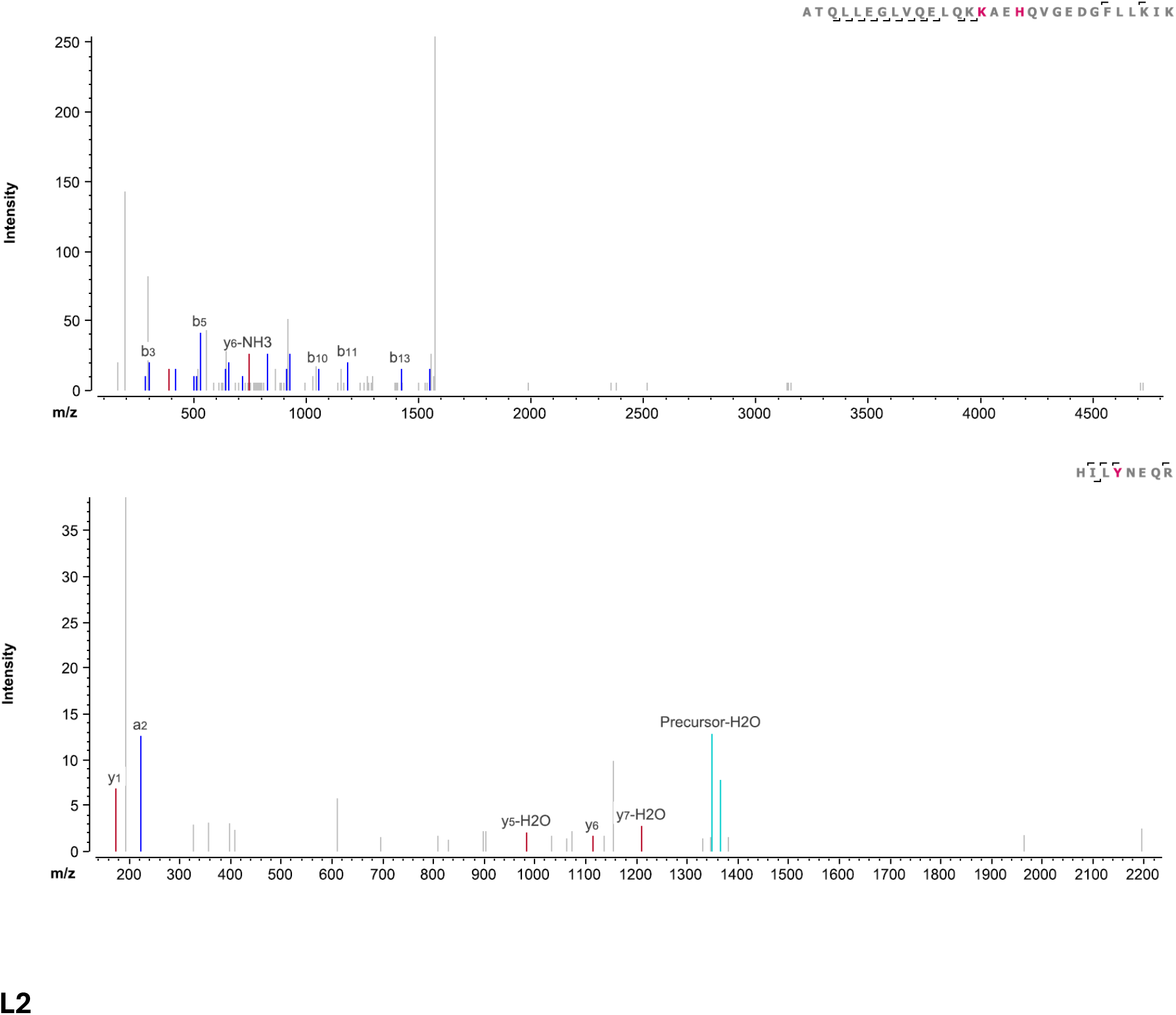
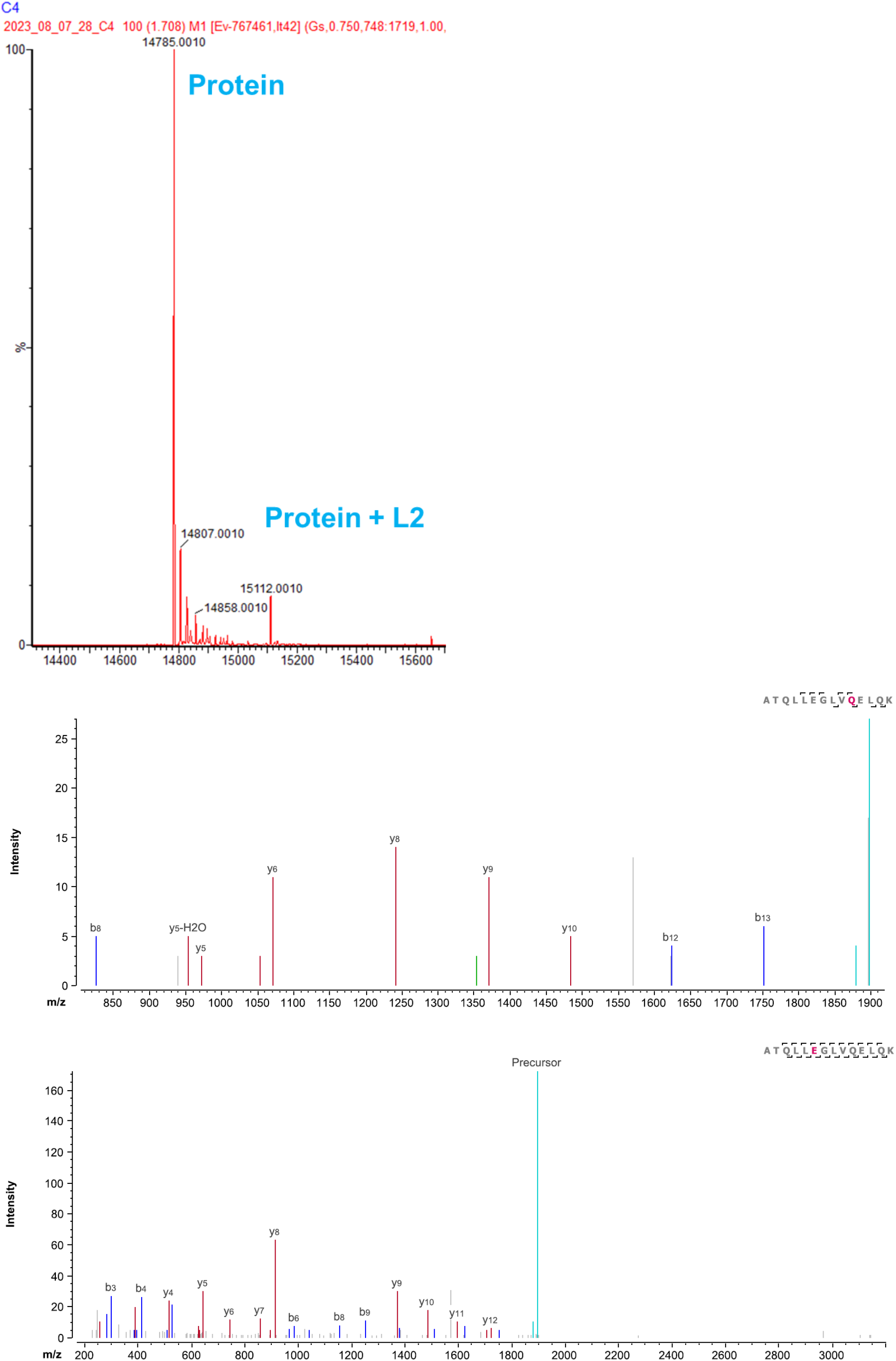
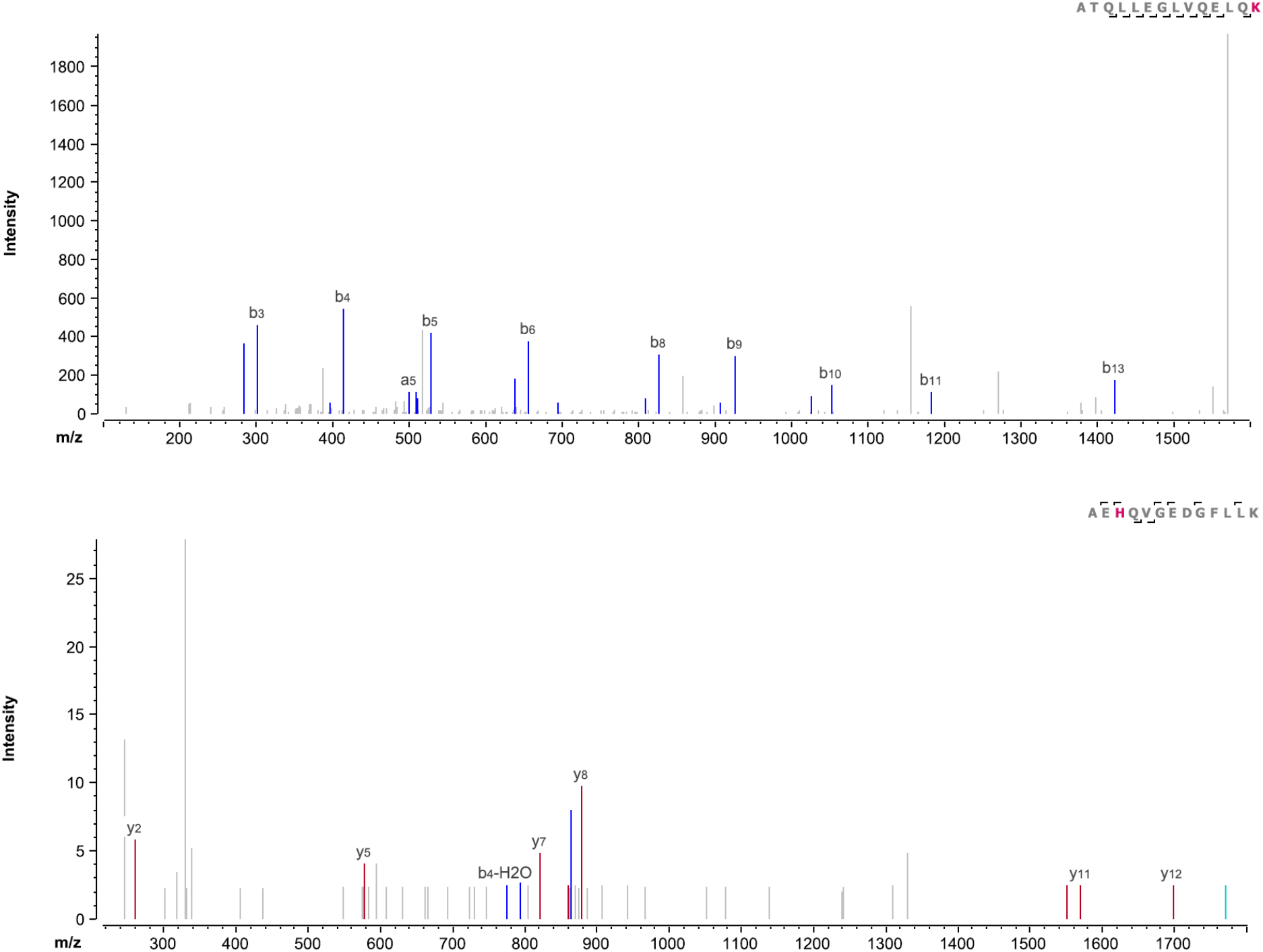

## 5. Characterization of fragment hits

Characterization of PhP060, PhP071, PhP072, F2 (61), F4 (63), F5 (PhP012), F6 (PhP098), F7 (PhP018), F8 (PhP048), F10 (60), L1 (PhP038), L2 (PhP035) were previously reported.^1,11^ Fragments F1, F3 and F9 were synthesized according to the procedure described in ref.^1^

**F1 (H1M)**

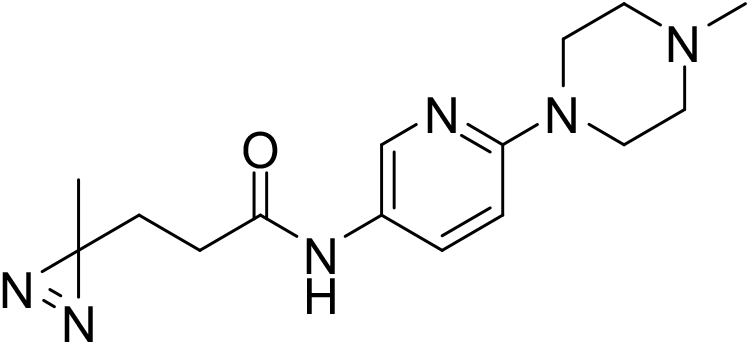

^1^H NMR (500 MHz, CDCl_3_) δ 8.17 (d, *J* = 2.5 Hz, 1H), 7.92 (dd, *J* = 9.1, 2.6 Hz, 1H), 7.46 (s, 1H), 6.65 (d, *J* = 9.1 Hz, 1H), 3.87 – 3.80 (m, 4H), 3.50 – 3.44 (m, 4H), 2.19 (t, *J* = 7.6 Hz, 2H), 1.84 (t, *J* = 7.6 Hz, 2H), 1.06 (s, 3H) ppm. ^13^C NMR (126 MHz, CDCl_3_) δ 170.13, 163.24, 156.55, 139.43, 132.13, 125.94, 107.29, 66.61, 46.08, 31.17, 29.89, 19.93 ppm.

Exact Mass [M+H]^+^: Calcd: 290.1538 Found: 290.1611

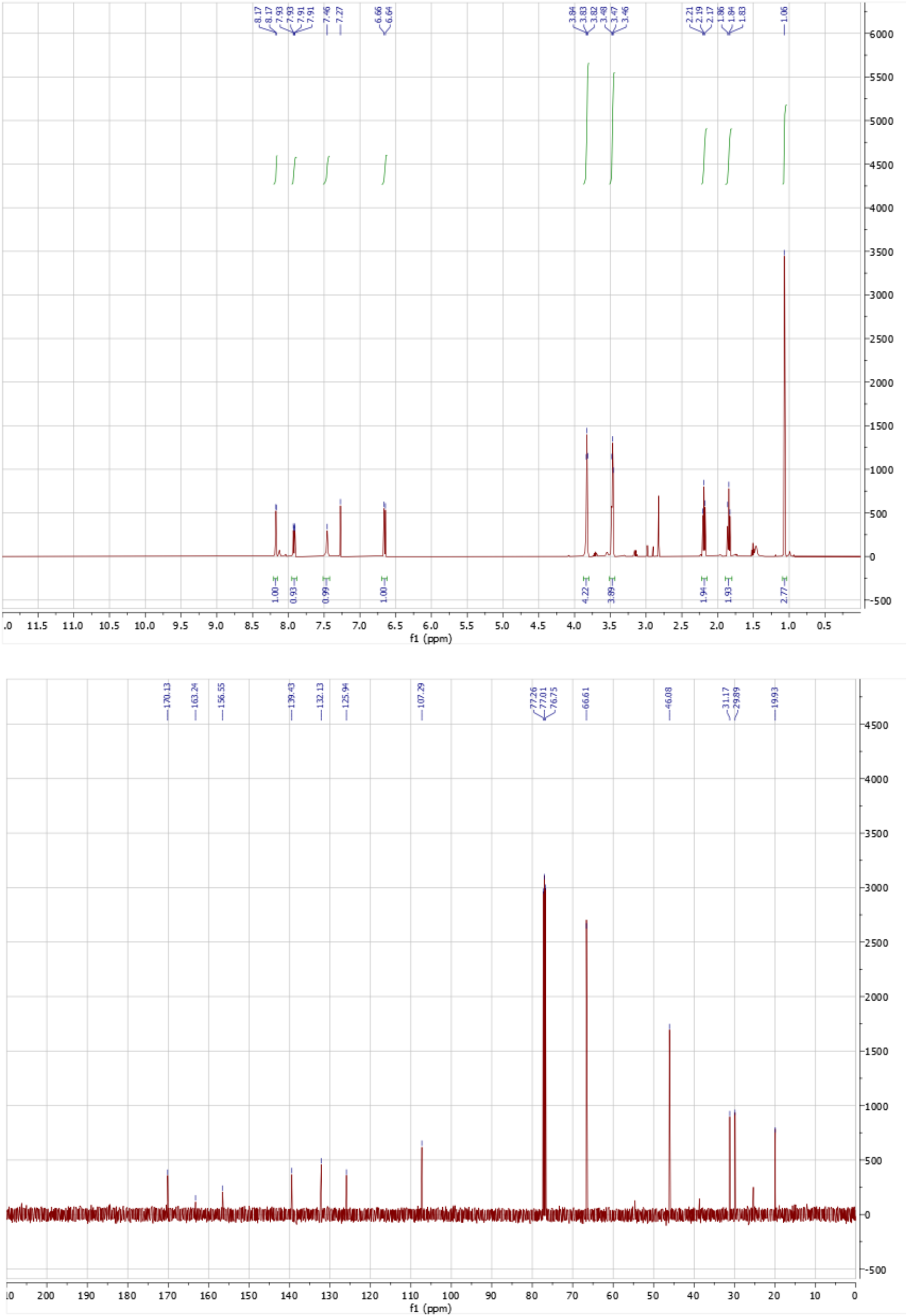

**F3 (H2M)**

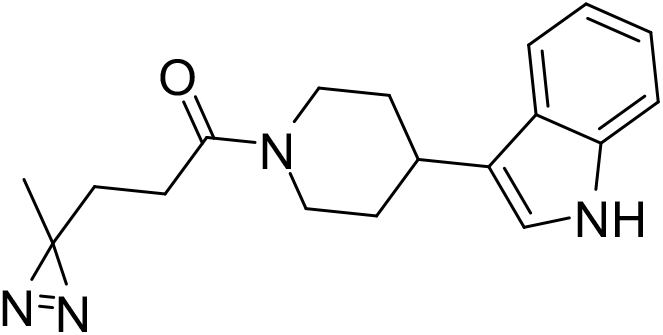

^1^H NMR (500 MHz, CD_3_CN) δ 9.10 (s, 1H), 7.66 (d, *J* = 8.0 Hz, 1H), 7.42 (d, *J* = 8.1 Hz, 1H), 7.19 – 7.13 (m, 1H), 7.10 – 7.03 (m, 2H), 4.64 (d, *J* = 13.1 Hz, 1H), 3.97 (d, *J* = 13.7 Hz, 1H), 3.32 – 3.18 (m, 1H), 3.17 – 3.06 (m, 1H), 2.85 – 2.70 (m, 1H), 2.37 – 2.24 (m, 2H), 2.07 (t, *J* = 14.4 Hz, 2H), 1.80 – 1.50 (m, 4H), 1.07 (s, 3H) ppm. ^13^C NMR (126 MHz, CD_3_CN) δ 169.39, 136.66, 126.51, 121.47, 120.35, 119.98, 117.28, 117.17, 111.40, 45.82, 42.11, 33.51, 33.26, 32.57, 29.69, 26.94, 19.01 ppm.

Exact Mass [M+H]^+^: Calcd: 311.1793 Found: 311.1866

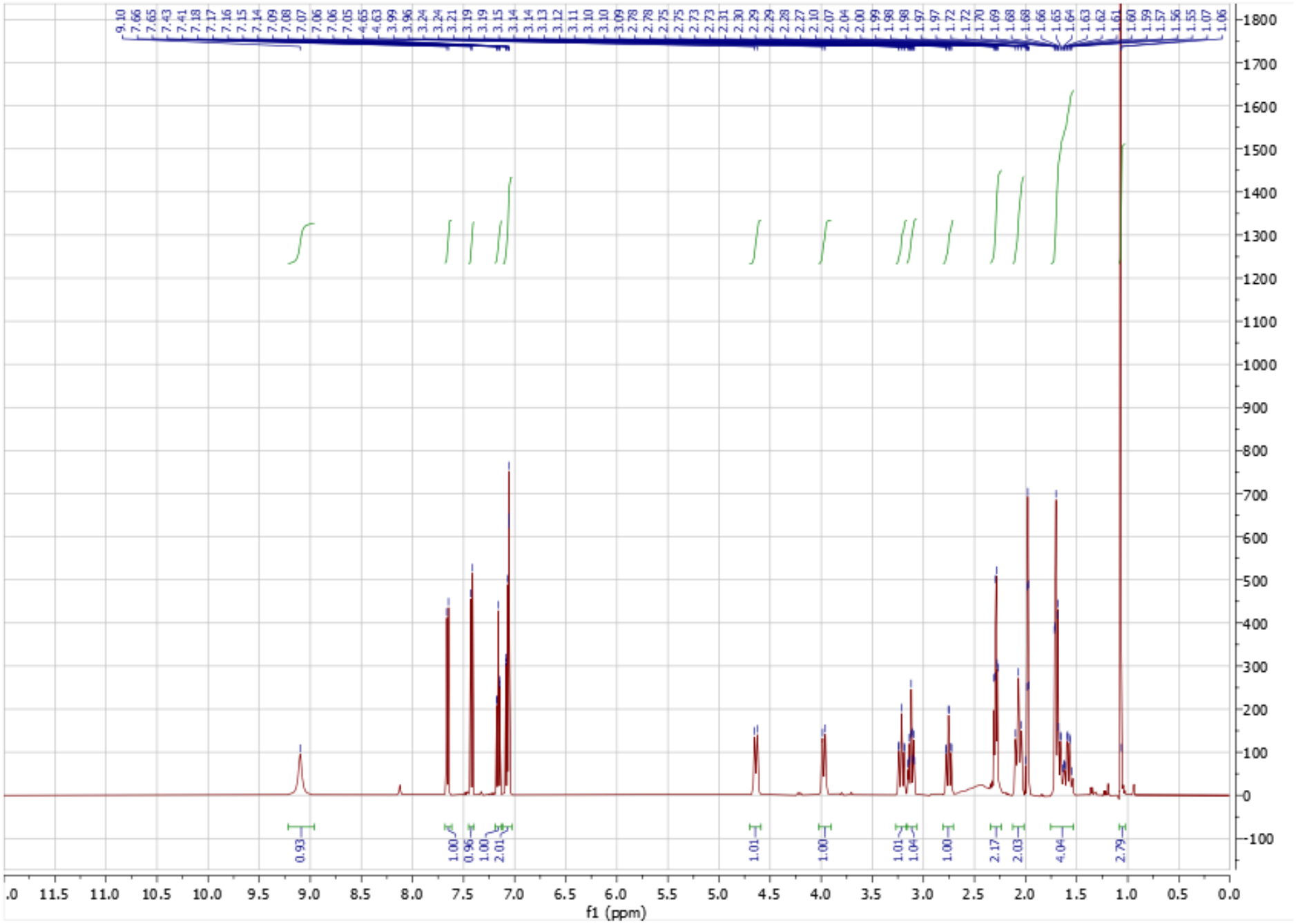
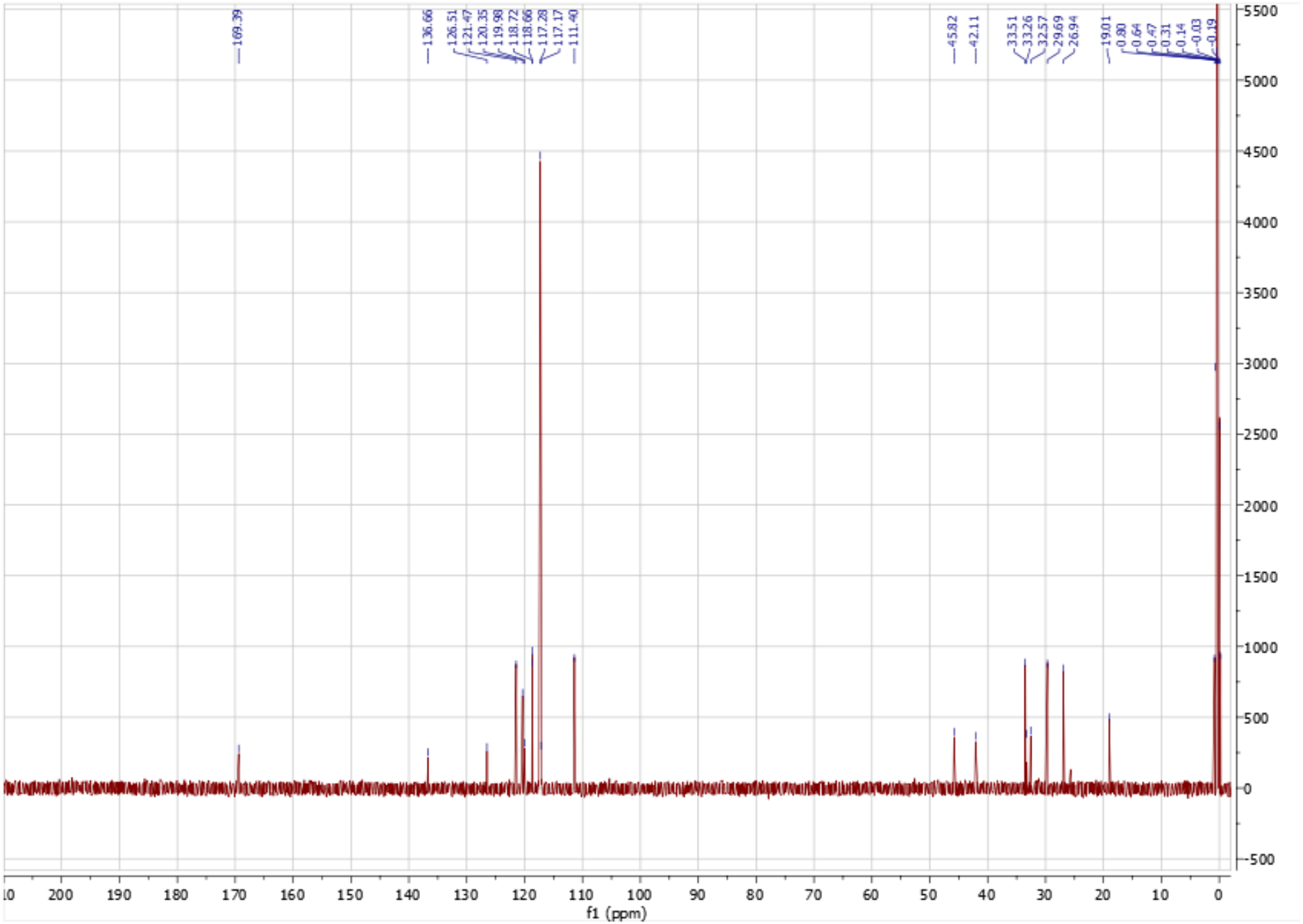

**F9 (H3M)**

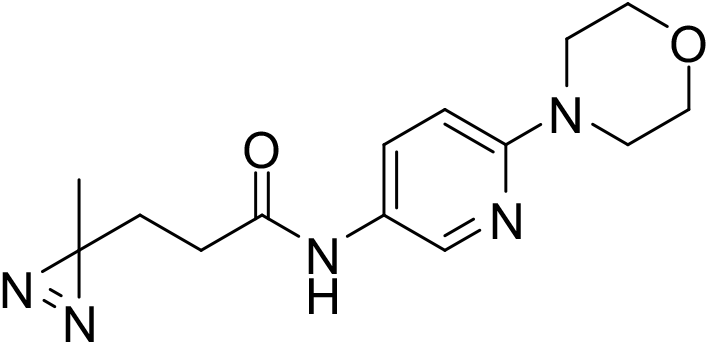

^1^H NMR (500 MHz, CD_3_CN) δ 8.31 (d, *J* = 2.5 Hz, 1H), 8.26 (d, *J* = 42.7 Hz, 1H), 7.82 (dd, *J* = 9.0, 2.6 Hz, 1H), 6.82 (d, *J* = 9.0 Hz, 1H), 3.82 – 3.67 (m, 2H), 3.30 (s, 2H), 2.87 (s, 2H), 2.79 (s, 3H), 2.25 (t, *J* = 7.6 Hz, 2H), 1.75 (t, *J* = 7.6 Hz, 2H), 1.35 (d, *J* = 6.6 Hz, 2H), 1.06 (s, 3H) ppm. ^13^C NMR (126 MHz, CD_3_CN) δ 170.43, 154.91, 139.89, 130.95, 127.75, 107.75, 53.19, 42.97, 42.85, 37.96, 30.63, 29.70, 25.54, 18.84 ppm.

Exact Mass [M+H]^+^: Calcd: 303.1855 Found: 303.19270485

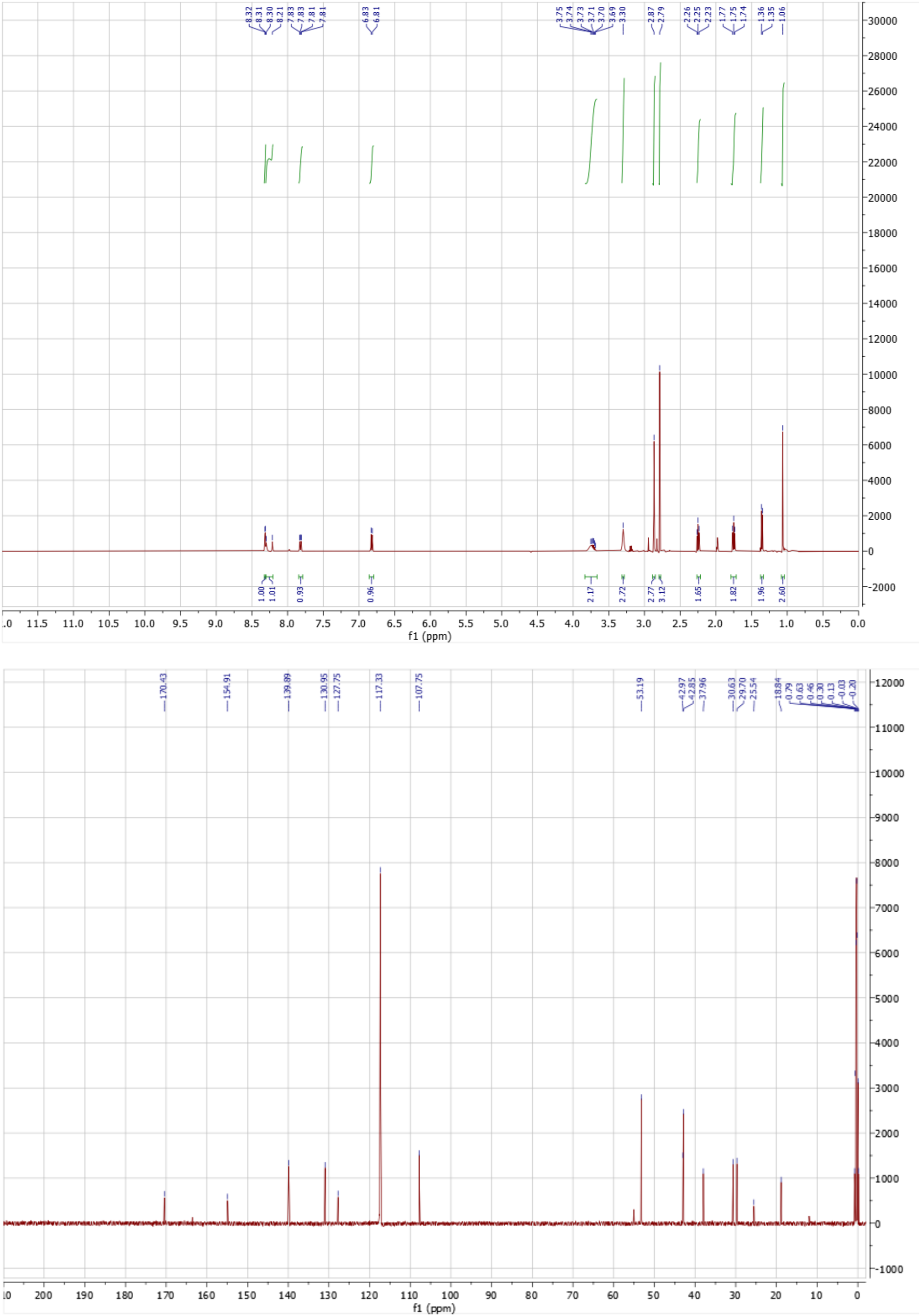

## Notes

https://doi.org/10.5281/zenodo.21266702

https://github.com/keserulab

### Competing Interest Statement

The authors have declared no competing interest.

